# Transcriptome Profiles of *Anopheles gambiae* Harboring Natural Low-Level Plasmodium Infection Reveal Adaptive Advantages for the Mosquito

**DOI:** 10.1101/2021.08.02.454766

**Authors:** Ann Carr, David C. Rinker, Yuemei Dong, George Dimopoulos, Laurence J. Zwiebel

**Affiliations:** Department of Biological Sciences, Vanderbilt University, Nashville, TN 37235, USA; Department of Chemistry, Vanderbilt University; W. Harry Feinstone Department of Molecular Microbiology and Immunology, Bloomberg School of Public Health, Johns Hopkins University, Baltimore, MD 21205, USA

## Abstract

Anopheline mosquitoes are the sole vectors for the *Plasmodium* pathogens responsible for malaria, which is among the oldest and most devastating of human diseases. The continuing global impact of malaria reflects the evolutionary success of a complex vector-pathogen relationship that accordingly has been the long-term focus of both debate and study. An open question in the biology of malaria transmission is the impact of naturally occurring low-level *Plasmodium* infections of the vector on the mosquito’s health and longevity as well as critical behaviors such as host-preference/seeking. To begin to answer this, we have completed a comparative RNAseq-based transcriptome profile study examining the effect of biologically salient, salivary gland transmission-stage *Plasmodium* infection on the molecular physiology of *Anopheles gambiae* s.s. head, sensory appendage, and salivary glands. When compared with their uninfected counterparts, *Plasmodium* infected mosquitoes exhibit increased transcript abundance of genes associated with olfactory acuity as well as a range of synergistic processes that align with increased fitness based on both anti-aging and reproductive advantages. Taken together, these data argue against the long-held paradigm that malaria infection is pathogenic for anophelines and, instead, suggests there are biological and evolutionary advantages for the mosquito that drive the preservation of its high vectorial capacity.

## Introduction

Globally, malaria remains the most endemic infectious/mosquito-borne disease, with over 500 million cases per annum, and putative vectors present in almost 100 countries worldwide placing up to 40% of the world’s population at risk (malERA, 2011). Human malaria is the result of pathogenic infection by five species of unicellular *Plasmodium Haemosporidians* (WHO, 2020). Of utmost importance is *Plasmodium falciparum*, which causes severe health complications and the highest human mortality rate (Olliaro, 2008). *Plasmodium* parasites are solely vectored between humans by mosquitoes within the genus *Anopheles* (Coetzee and Koekemoer, 2013; Sinka et al., 2012). *Anopheles* mosquitoes are anautogenous; females require a blood meal for reproduction and replenishment of energy stores (Hagan (2018). Importantly, *An. gambiae* adult females are only infectious to humans when the *Plasmodia* they harbor complete a multi-stage sporogonic cycle and successfully invade the mosquito’s salivary glands. Despite the complexity of the interlaced *Anopheles-Plasmodium* life cycles and the inherent ecological and environmental challenges, it is often surprising that in malaria-endemic regions of Sub-Saharan Africa, where there is a high disease prevalence, the percentage of infectious mosquitoes within surveys of *Anopheles* populations is actually very low, typically ranging from 1% to 10% (Ogola et al., 2018). Blood feeding, which is carried out solely by adult females, is a central factor that positively affects the complex *Anopheles-Plasmodium* paradigm. As such, *Anopheles* blood-meal host-seeking and preference are critical behaviors for regulating the synergistic processes that ultimately impact the mosquito’s vectorial capacity. Indeed, several studies support a hypothesis that pathogenic malaria in humans and *Plasmodium* infection of vectors may modulate mosquito behavior and physiology to increase the likelihood of pathogen transmission (Anderson et al., 1999; Lacroix et al., 2005; Stanczyk et al., 2019; Stanczyk et al., 2017). However, many of these and other efforts to examine the relationship between *Anopheles and Plasmodium* have for experimental simplicity often resorted to employing unnaturally high mosquito infection intensities that might have artifactually altered the natural relationships between *Anopheles* and *Plasmodium* (Ferguson and Read, 2004; Pinheiro-Silva et al., 2015; Smallegange et al., 2013; Stanczyk et al., 2017).

To begin, the vast majority of anopheline mosquitoes that ingest *Plasmodium* gametocyte-infected blood meals never become infectious (Belachew, 2018). In controlled laboratory studies, only 5-10% of ingested *Plasmodium* gametocytes successfully differentiate into ookinetes; of those, less than 0.3% form oocysts, giving rise to approximately 1-2 oocysts per mosquito midgut (Belachew, 2018; Vaughan et al., 1994). This closely mirrors West African field studies indicating that the majority of malaria-infected mosquitoes are within that range and only rarely are trapped mosquitoes observed with more than 5 oocysts per midgut (Collins and Besansky, 1994). In that light, it is reasonable to suggest that when investigating salient biological relationships between *Anopheles* and *Plasmodium* in the laboratory, the relative intensity of mosquito infections should align with these natural levels. The high malaria infection intensities that are often employed in studies to provide high infection prevalence are likely to significantly alter the *Anopheles-Plasmodium* paradigm potentially giving rise to potentially biologically irrelevant shifts in mosquito gene expression, physiology and behavior. We have deliberately employed low gametocytemia and, thereby, biologically relevant *P. falciparum* blood meals to examine how salivary-gland-stage sporozoite infection influences the transcriptome of female *An. gambiae* head (including sensory appendages) and salivary glands, henceforth referred to as ‘head’. These data suggest that natural levels of malaria infection align with enhanced olfactory sensitivity that impacts chemosensory driven host preference. Moreover, and surprisingly, natural levels of *Plasmodium* infection appear to provide a broad and synergistically beneficial modulation of mosquito fitness that underlies and may indeed promote the evolutionary stability of the *Anopheles-Plasmodium* paradigm that continues to drive global malaria transmission.

## Results and Discussion

### Experimental Design & Establishment of Optimal Gametocytemia

Our study was designed to rigorously compare the impact of biologically salient *P. falciparum* infections on the *An. gambiae* head (encompassing brain, sensory appendages, and salivary glands) transcriptome. An experimental paradigm incorporating triplicate biological replicates (21 samples per replicate) was designed wherein 5-day-old females were blood fed on either *P. falciparum*-infected (treatment) or naïve (control) human blood meals and thereafter reared for 18 days prior to collection and dissection for downstream molecular analyses. This paradigm allows for completion of the mosquito sporogonic cycle as well as sporozoite invasion of the salivary glands in treated (infected) mosquitoes (Figure 1A), thus ensuring that mosquitoes harboring sporozoites in their salivary glands are examined. Within this study, the importance of establishing and utilizing the low-level intensities representative of natural biological infections was prioritized, in spite of the experimental challenges resulting from significant reductions in prevalence (Collins and Besansky, 1994). To achieve this goal, approximately 14% of the treatment mosquitoes were sampled 8 days after blood meals, and their midguts were examined for the presence of *P. falciparum* oocysts. Preliminary studies indicated that mosquitoes that had blood fed on human blood supplemented with *P. falciparum* NF54W at a final gametocytemia of 0.15%, 0.05%, and 0.01% resulted in median/upper threshold oocyst counts that were significantly higher than the optimal levels of 1-oocyst/midgut median and 3-6 oocysts/midgut upper levels required. Iterative efforts established an optimal gametocyte dilution of 0.008% to meet those criteria, which was thereafter used for all treatments in this study (Table 1). The heads (including sensory appendages and salivary glands) were used for preparation of total RNA (for sequencing) and salivary tissues for genomic DNA (gDNA) for PCR validation of the presence of *P. falciparum* sporozoite-treated individual mosquitoes. These PCR-based analyses utilized primers designed to detect the *P. falciparum* circumsporozoite (CS) gene as well as the *An. gambiae* ribosomal protein s7 (AgRsp7) gene as an internal standard (Coppi et al., 2011). This analysis revealed that, of the 176 treatment mosquitoes that took infected blood meals and survived to 23 days of age, 65 were confirmed as having salivary-gland-stage sporozoites, representing an overall prevalence of ∼37% (Figure 2). Consistent survivor rates of ∼50% were observed between treatment and control *An. gambiae* lines. The observed survival rate parallels previous studies documenting 20- day-old *An. gambiae* laboratory survival rates (Huestis et al., 2017).

**Figure 1.**
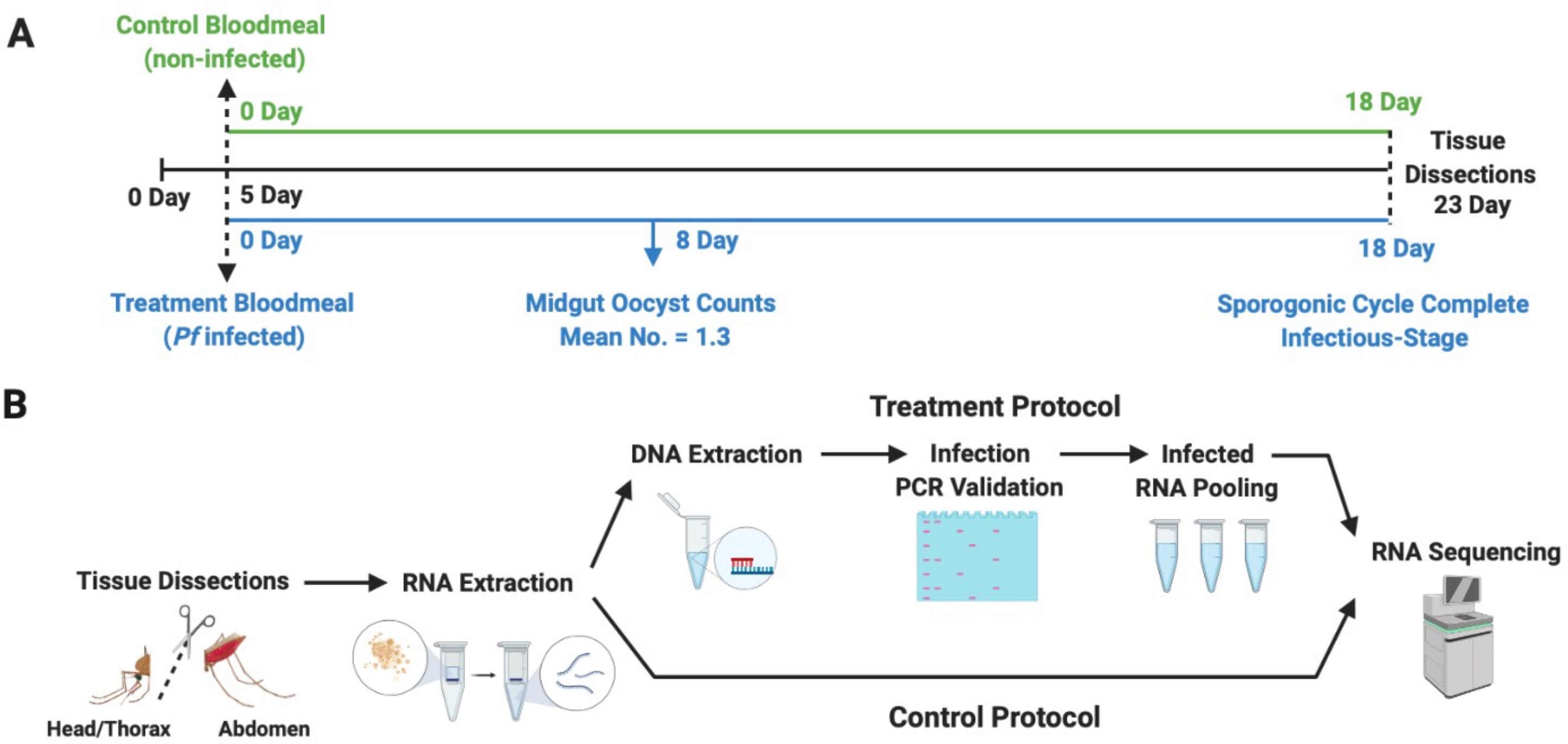
Schematic representation of experimental methodology. (A) Timeline for establishment of *Plasmodium falciparum* sporozoite infected treatment and naive controls *An. gambiae* s.s. mosquito lines. (B) Total RNA and gDNA extraction treatment and control protocols for PCR *P. falciparum* sporozoite infection validation and RNA sequencing.

**Figure 2.**
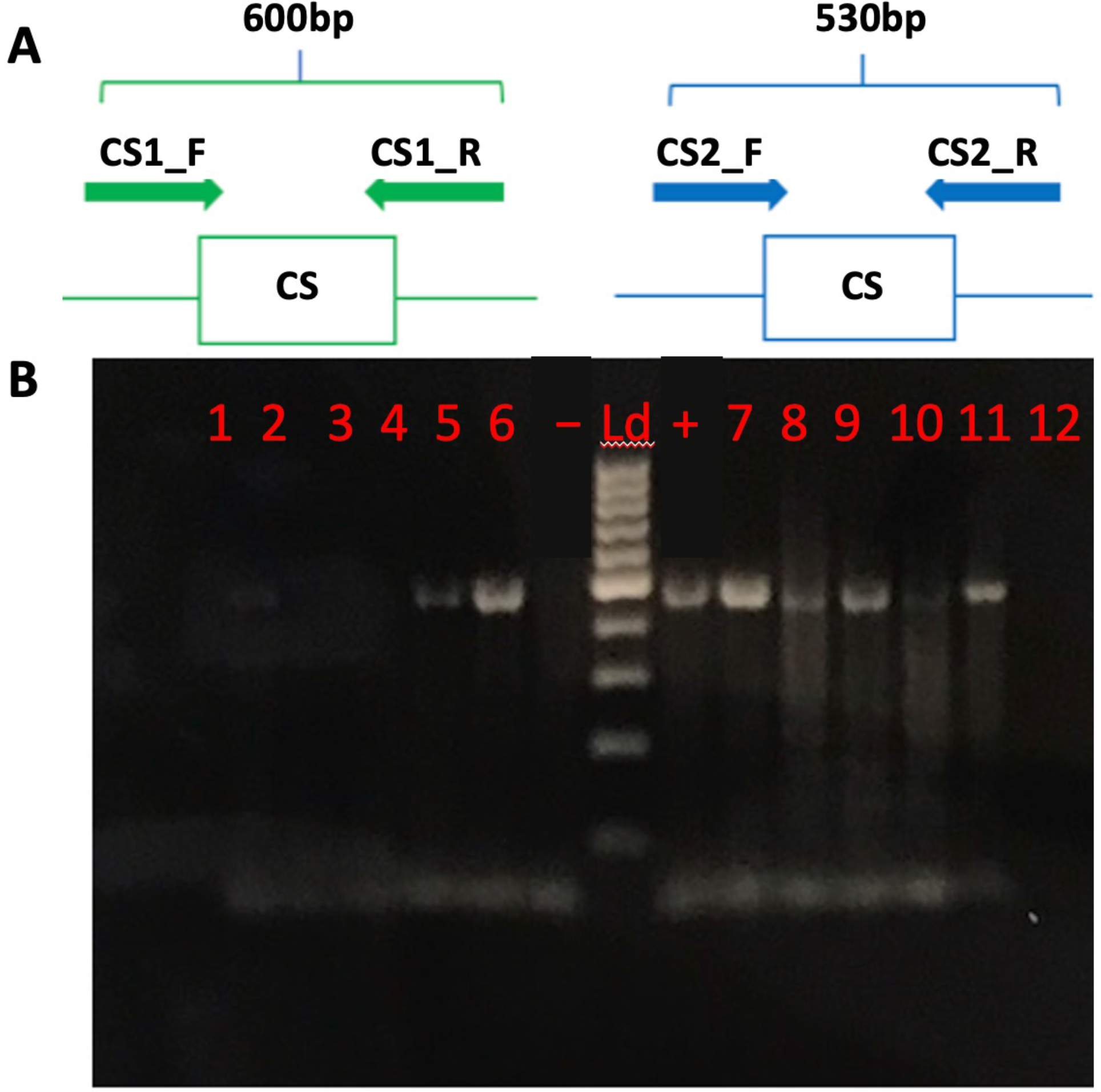
(A) *P. falciparum* circumsporozoite (CS) primer set schematics. (B) Agarose gel (1.5%) electrophoresis image of amplified products using CS2 primer sets. Lanes 1-6 and 10-16 are examined *P. falciparum* sporozoite infected treatment *An. gambiae* s.s. mosquitoes with a (Kazuya Ujihara) positive and (-) negative control. Lane Ld, 100bp DNA size marker.

**Table 1.**
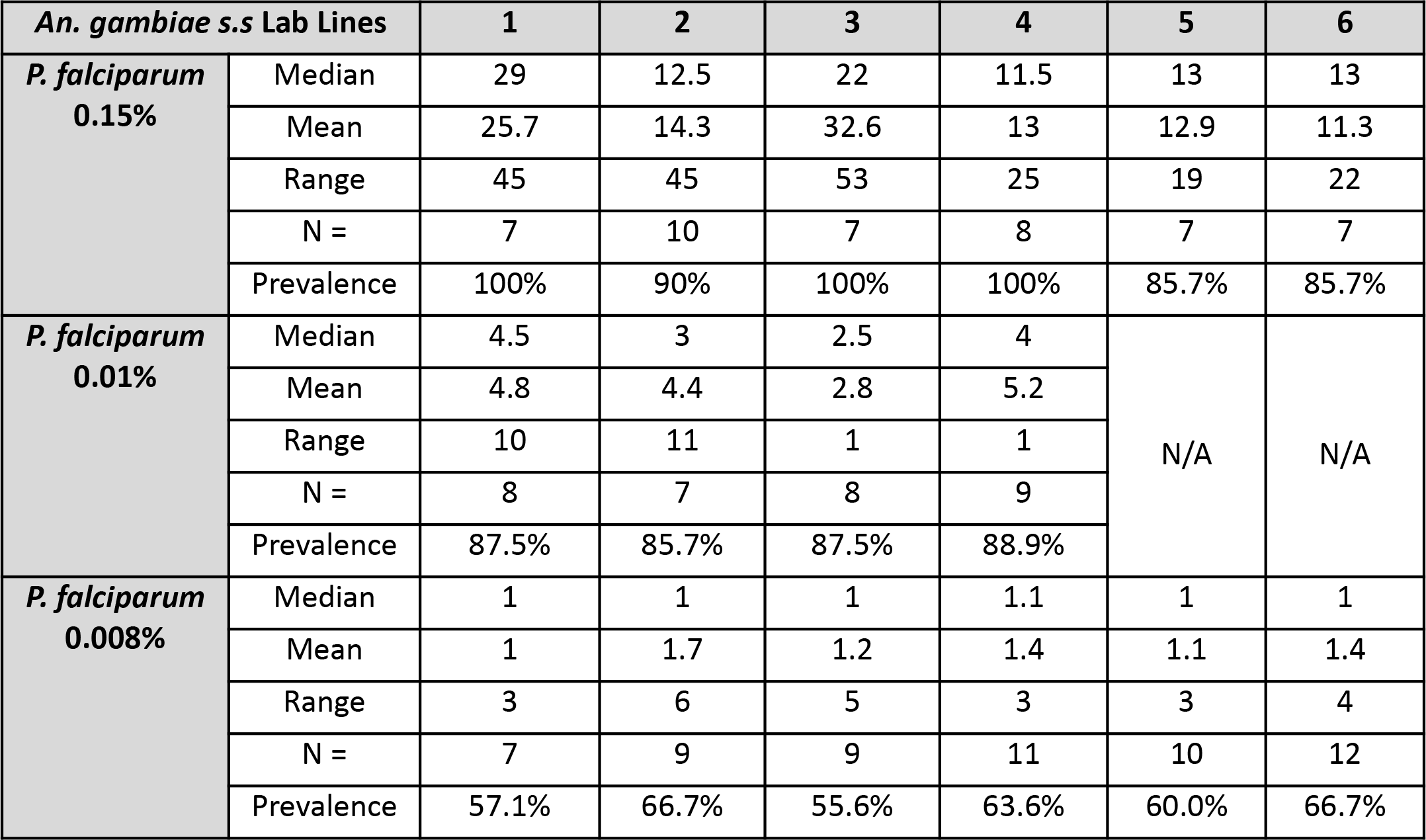
Results from midgut oocyst counts of 8-d.o. *P. falciparum* infected treatment *An. gambiae* s.s. mosquitoes.

| **An. gambiae s.s Lab Lines** |  | **1** | **2** | **3** | **4** | **5** | **6** |
| --- | --- | --- | --- | --- | --- | --- | --- |
| **P. falciparum**<br>**0.15%** | Median | 29 | 12.5 | 22 | 11.5 | 13 | 13 |
|  | Mean | 25.7 | 14.3 | 32.6 | 13 | 12.9 | 11.3 |
|  | Range | 45 | 45 | 53 | 25 | 19 | 22 |
|  | N = | 7 | 10 | 7 | 8 | 7 | 7 |
|  | Prevalence | 100% | 90% | 100% | 100% | 85.7% | 85.7% |
| **P. falciparum**<br>**0.01%** | Median | 4.5 | 3 | 2.5 | 4 | N/A | N/A |
|  | Mean | 4.8 | 4.4 | 2.8 | 5.2 |  |  |
|  | Range | 10 | 11 | 1 | 1 |  |  |
|  | N = | 8 | 7 | 8 | 9 |  |  |
|  | Prevalence | 87.5% | 85.7% | 87.5% | 88.9% |  |  |
| **P. falciparum**<br>**0.008%** | Median | 1 | 1 | 1 | 1.1 | 1 | 1 |
|  | Mean | 1 | 1.7 | 1.2 | 1.4 | 1.1 | 1.4 |
|  | Range | 3 | 6 | 5 | 3 | 3 | 4 |
|  | N = | 7 | 9 | 9 | 11 | 10 | 12 |
|  | Prevalence | 57.1% | 66.7% | 55.6% | 63.6% | 60.0% | 66.7% |

### RNAseq/Bioinformatics

Transcriptome profiling was conducted using total RNA extracted from triplicate *An. gambiae* treatment (infected; T1-3) and control (uninfected; C1-3) heads. Following sequence trimming and quality control bioinformatic pipelines, individual data sets were aligned to the *An. gambiae* PEST strain genome. Sequencing of the T1, T2, and T3 *P. falciparum*-infected (treatment) *An. gambiae* head libraries generated a total of over 254 million reads mapping to 83%, 83%, and 85% respectively of the *An. gambiae* genome (VectorBase-53_AgambiaePEST; Table 2). Likewise, Illumina sequencing of the C1, C2, and C3 control (uninfected) *An. gambiae* head libraries generated a total of 426 million reads mapping to 81%, 80%, and 81% respectively of the *An. gambiae* genome (VectorBase-53_AgambiaePEST; Table 2). Uniquely mapped reads were counted using the program HTSeq. DESeq2 statistical analysis identified 5,765 annotated transcripts present in significantly different abundances between treatment and control replicates (*p* < 0.05; Figure 3A). These annotated transcripts represent a transcriptional shift of 36% of the *An. gambiae* head transcriptome. Of these, 2,776 transcripts were present in significantly higher abundances in treatment replicates and the remaining 2,989 transcripts present in significantly higher abundance in control replicates (*p* < 0.05). With such a large transcriptional shift observed between treatment and control *An. gambiae* heads, both principal component analysis (PCA) and Pearson’s correlation coefficient (PCC) calculations were performed to verify validity. PCC calculations for the three treatment and three control *An. gambiae* head libraries returned *r* values of 0.9429 and 0.9770, respectively, indicating a strong association among biological replicates (Table 2). Additionally, PCA determined that 83% of observed variance between the six library datasets was attributed to the presence/absence of *P. falciparum* infection, with negligible variance attributed to biological replication (Figure 3B).

**Figure 3.**
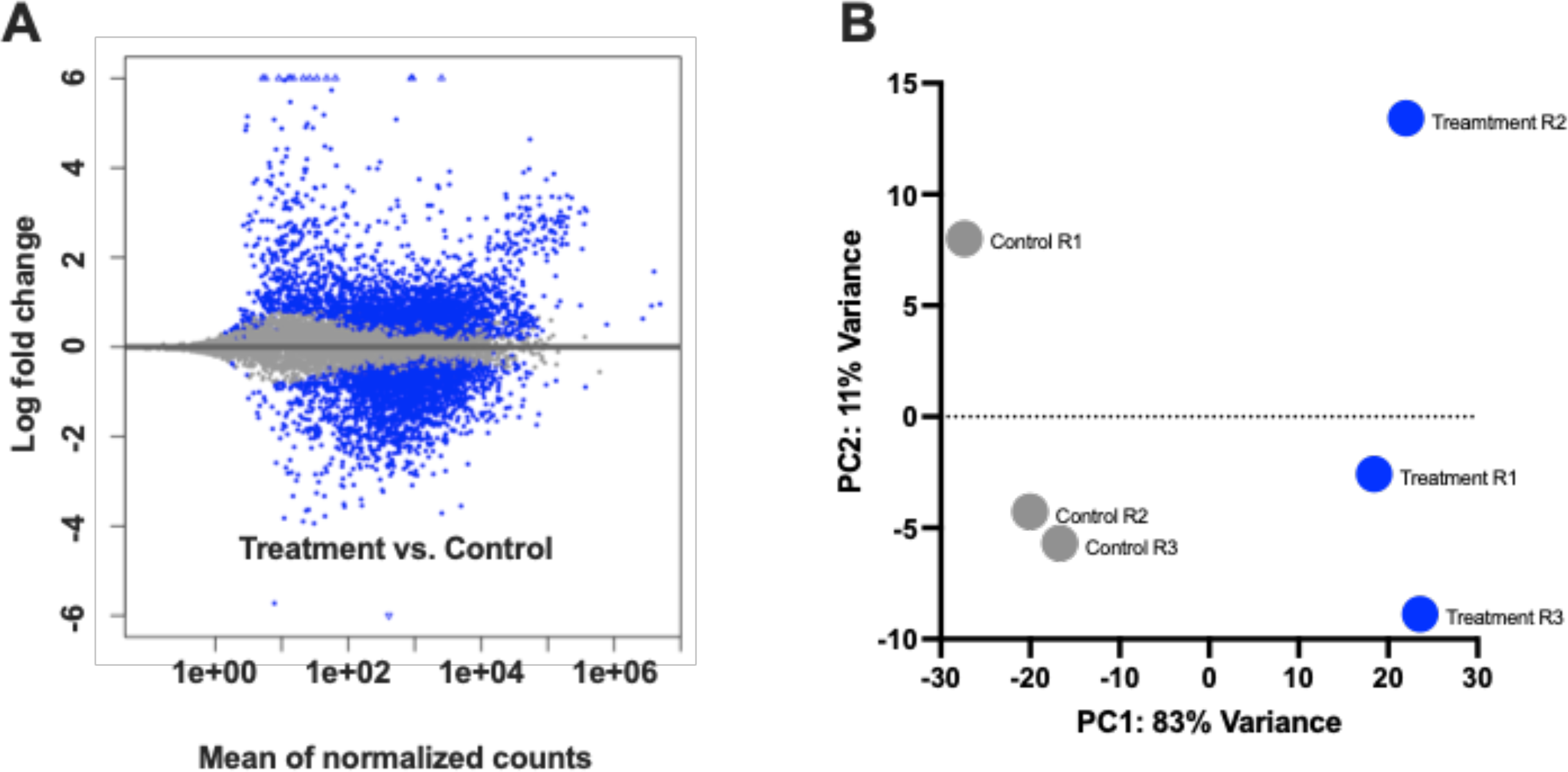
(A) Scatterplot of log fold changes vs. mean normalized counts generated using HTSeq and DESeq2 for *P. falciparum* sporozoite infected treatment and naïve control library replicates (p < 0.05). (B) Principal component analysis of the 3 sporozoite infected treatment and 3 naïve control library replicates. Gray circles represent control replicates; blue circles represent treatment replicates.

**Table 2.** Statistics of RNA sequencing for *P. falciparum* sporozoite infected treatment and naïve control *An. gambiae* s.s. libraries.

| Exp. Line | Replicate | Total No. Reads | Mapped Reads | No. Transcripts | PCC (r) | Coverage |
| --- | --- | --- | --- | --- | --- | --- |
| Treatment | T1 | 65,613,838 | 54,459,486 (83%) | 12,194 (77%) | 0.9429 | 70x |
|  | T2 | 123,249,032 | 102,296,697 (83%) | 12,799 (81%) |  | 131x |
|  | T3 | 65,668,388 | 55,818,130 (85%) | 12,533 (79%) |  | 70x |
| Control | C1 | 166,457,374 | 123,178,457 (74%) | 12,848 (81%) | 0.9770 | 177x |
|  | C2 | 110,612,110 | 81,852,961 (74%) | 12,625 (80%) |  | 118x |
|  | C3 | 149,580,984 | 107,698,308 (72%) | 12,719 (81%) |  | 160x |

GO annotation using OmicsBox assigned 4,976 GO-terms (Level II) to 2,832 transcripts of the 5,765 differentially abundant transcripts (control and treatment libraries). Of the total annotated transcripts, 1,291 were elevated in treatment libraries and 1,541 transcripts elevated in control libraries, with approximately 10% of the transcripts having no known function or characterization (Figure 4).

**Figure 4.**
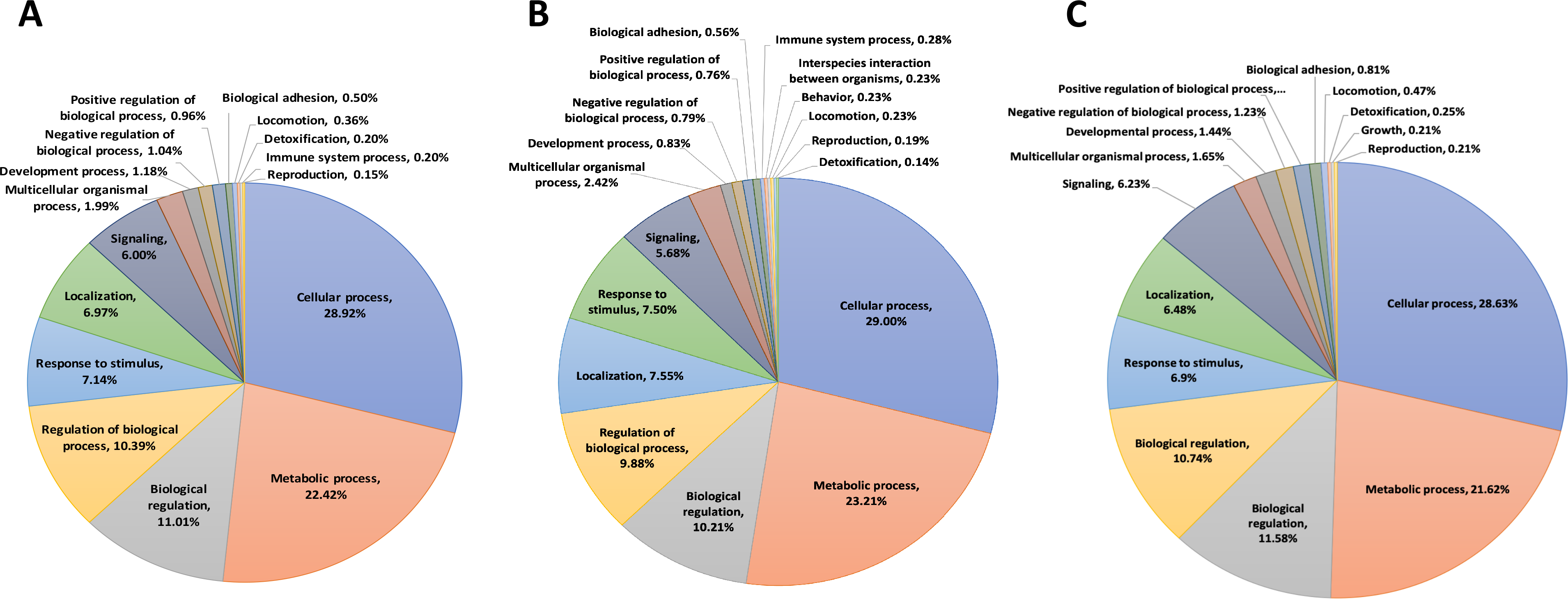
Distribution of transcripts annotated at the gene ontology level 2 and their putative involved in biological functions for (A) all transcripts differentially abundant between *P. falciparum* sporozoite infected treatment *An. gambiae* s.s. and naïve controls, (B) only transcripts differentially abundant in sporozoite infected treatments, and (C) only transcripts differentially abundant in naïve controls.

Transcripts with no clear molecular function via GO were further BLASTed using Blast2GO and the GeneBank non-redundant protein database to assist in further functional analysis. The resultant assigned GO annotations are representative of a diverse array of biological processes and molecular functions that make interpretation difficult. Taken together, these analyses reveal a broad transcriptome profile shift that spans a wide range of cellular processes, some of which are likely to impact the behavior, physiology and ultimately the vectorial capacity of *An. gambiae* carrying biologically relevant *P. falciparum* sporozoite infections.

### Plasmodium Manipulation of Mosquito Chemosensory Mechanisms

To examine the potential for behavioral modification of infected mosquitoes, our initial focus was to examine transcriptome modulations directly relevant to the chemosensory systems of *An. gambiae*. These systems collectively play an essential role in establishing and maintaining critically important behaviors, including the blood-meal host preference of female mosquitoes (Montell and Zwiebel, 2016; Takken and Verhulst, 2013).

### Odorant Receptors

The transcript abundance of several *An. gambiae* odorant receptors (AgORs) were highly differentiated between *P. falciparum* sporozoite-infected (sporozoite; treatment) and uninfected (control) heads. Indeed, 60 of the 71 AgORs and the OR co-receptor (Orco) that were detected in sequencing libraries were consistently more abundant in heads of infected mosquitoes, with 19 displaying statistically significant induction (Figure 5A). Transcripts encoding the remaining 11 AgORs (AgOR1, AgOR3, AgOR4, AgOR26, AgOR36, AgOR45, AgOR46, AgOR52, AgOR60, AgOR62 and AgOR72) showed an insignificant trend of less abundance in samples from infected mosquitoes (Figure 5A). With the near complete representation of AgORs (95% of annotated *An. gambiae* ORs), it is evident that sporozoite infection significantly impacts the levels of these ORs within mosquito chemosensory structures. The majority of anopheline ORs are functionally characterized, and we have previously integrated OR functional RNAseq data to model the receptivity profile for the antennae of *An. gambiae* following their initial bloodmeal (Rinker et al., 2013a).

**Figure 5.**
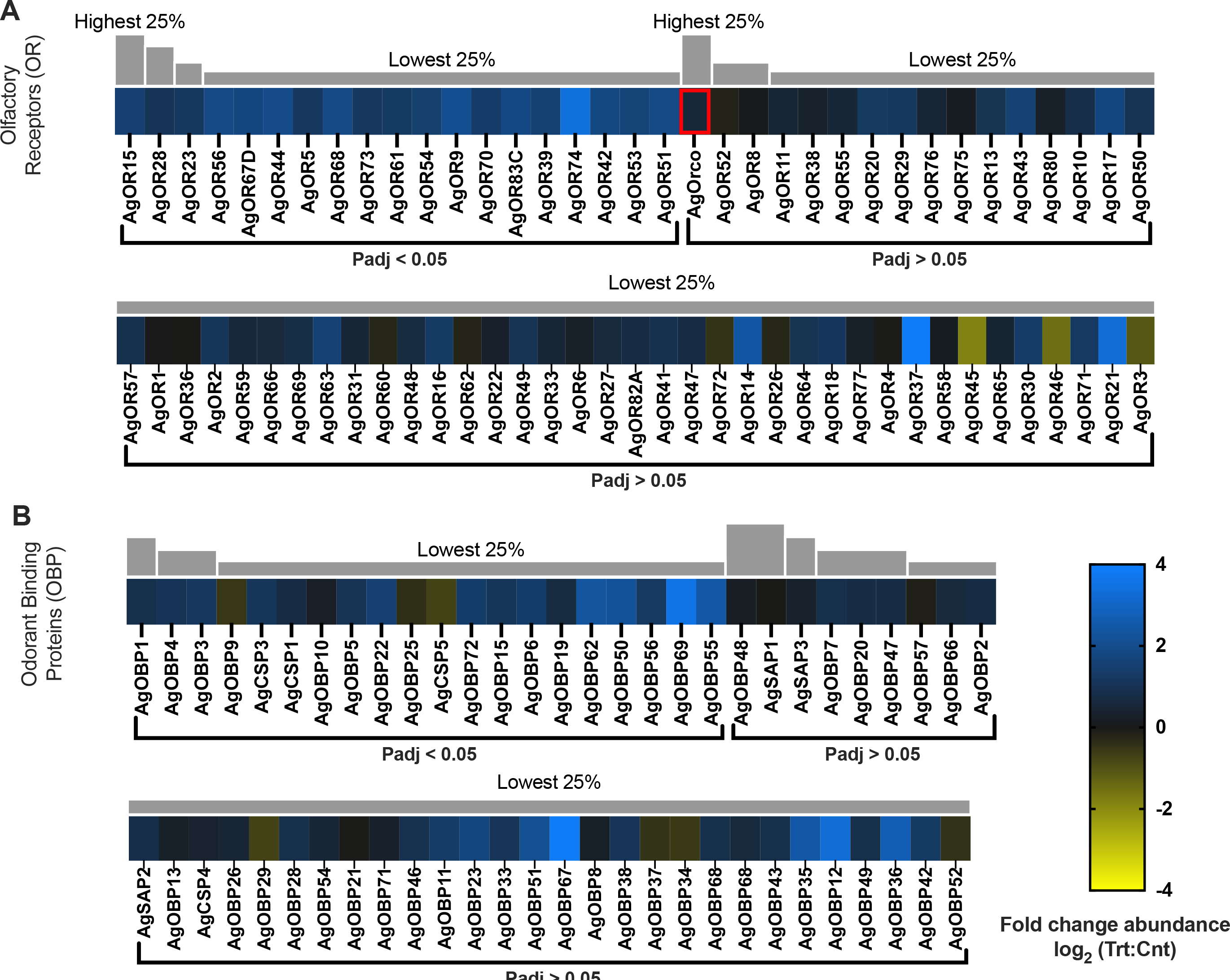
Chemosensory differential transcript abundances following relevant sporozoite infections. Chemosensory transcripts that were present at significantly higher (blue) or lower (yellow) levels in *P. falciparum* sporozoite infected treatments; non-differentially expressed chemosensory transcripts are denoted as zeros (black). Chemosensory genes within each family organized by adjusted p-value and subsequently arrayed left to right from most abundant to least abundant based on FPKM values (quartile bars above each image). (A) Odorant receptor family (OR). (B) Odorant binding protein family (Pattanakitsakul et al.). Chemosensory co-receptors are identified by red boxes. Log2 scale indicates transcript abundances that were significantly higher (blue) or lower (yellow) in sporozoite infected treatments (Vrba et al.) vs. naïve controls (Cnt).

Transcriptome profiling studies together with behavioral analyses support the hypothesis that the sensitivity of the anopheline olfactory system is directly associated with the relative abundance of olfactory transcripts (Rinker et al., 2013a; Rinker et al., 2013b; Wolff and Riffell, 2018). Applying this approach here suggests that the large number of significantly abundant AgOR transcripts associated with infected *An. gambiae* indicates substantial increases in odorant receptivity (Figure 6). In fact, AgOR receptivity modelling identified 70 odorants, many of which are components of human sweat, with significantly increased receptivity in *Plasmodium*-infected mosquitoes than in uninfected controls. Of these, several [including L-lactic-acid, 1-octen-3-ol and 4-methylphenol] are human sweat components that have been shown to be attractive kairomones for host-seeking female anophelines (Cork and Park, 1996; Smallegange et al., 2005). Notably, lactic acid showed a 217% increase in AgOR receptivity in treatments. Furthermore, the significant increases in receptivity to 1-butanol (247%), 3-methyl-1-butanol (260%), and 2,3-butanedione (252%) align with laboratory olfactometry and semi-field studies, indicating that these compounds significantly increase the number of anopheline catches when added to MMX traps baited with CO2 and a basic blend of amine/carboxylic acid attractants (Verhulst et al., 2011).

**Figure 6.**
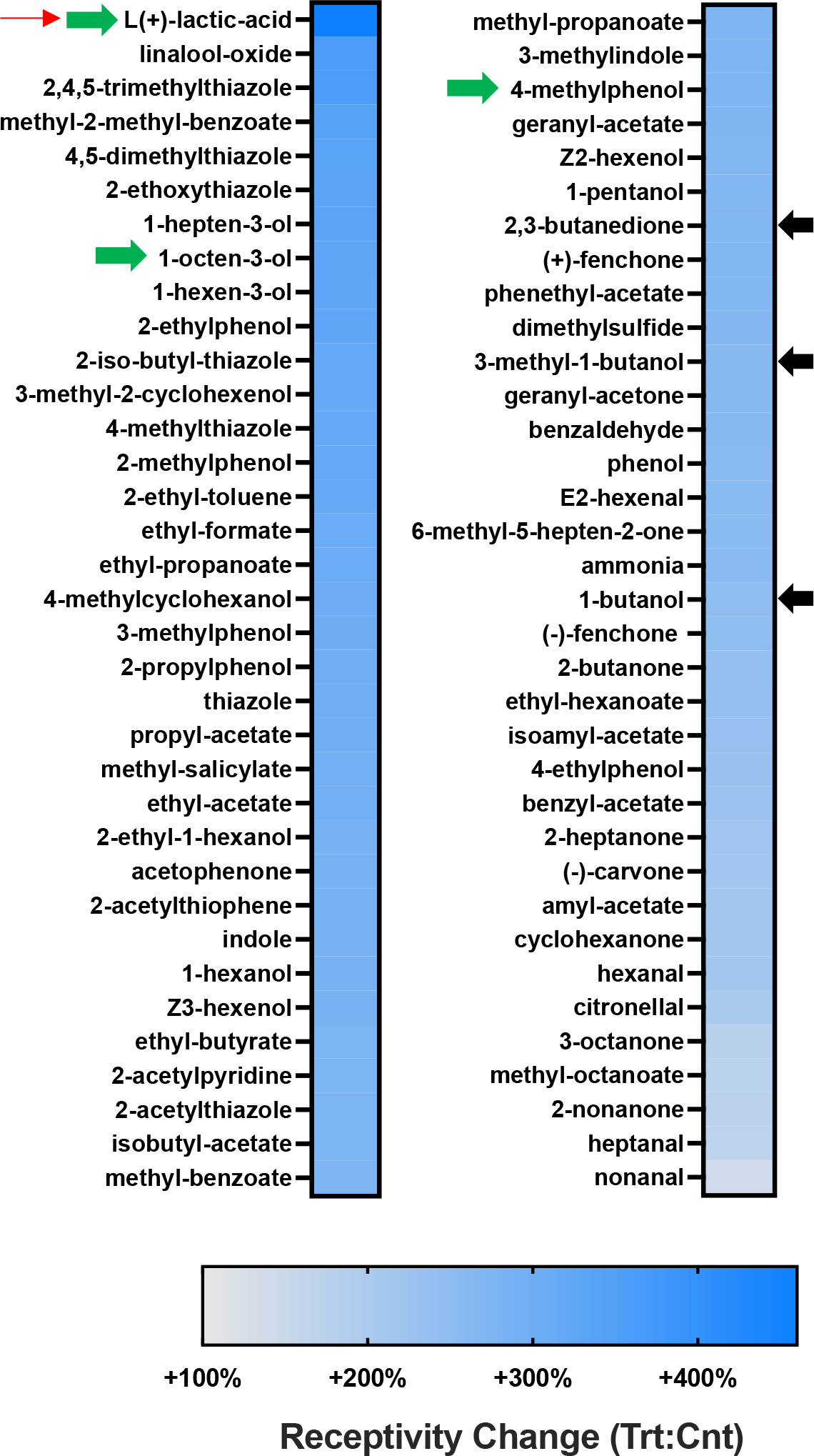
Calculated changes in AgOR mediated odorant receptivity following relevant sporozoite infections. Graphic representation of conceptualized differences in odorant receptivity for 70 odors in *P. falciparum* sporozoite infected treatments (Vrba *et al*.) vs. naïve controls (Cnt). Response characteristics were determined using known *An. gambiae* odorant receptor responses in heterologous expression systems and weighted by relative AgOR expression levels for each corresponding receptor. Results are sorted high to low. Scale bar show calculated increases (blue).

These putative increases of olfactory acuity in *Plasmodium*-infected anopheline mosquitoes are consistent with behavioral bioassay and surveillance studies that determined *Plasmodium* infections to increase mosquito bite frequency while maintaining host-preference plasticity (Cator et al., 2014; Vantaux et al., 2015). That said, our receptivity modelling does not align with recent studies that examined peripheral olfactory responses in *P. falciparum-*infected *An. gambiae* (Stanczyk et al., 2019). It is likely that the significantly higher infection intensities used in that study over-challenged *An. gambiae* infection-responsive systems, such as the innate immune system, resulting in unintended fitness consequences that altered host-seeking and blood-feeding behaviors (Ohm et al., 2016). Because our studies avoid impacts arising from non-natural infection intensities, they can more confidently reveal a clear effect of biologically relevant malaria infections on the anopheline olfactory system. The effect of increasing host seeking and host feeding synergistically support both parasite and mosquito survivorship as well as maximizing vectorial capacity. Interestingly, the presence of polycistronic OR genes in Anopheles (Karner et al., 2015) raises the hypothesis that *Plasmodium* parasites may have evolved to specifically activate the transcription factors regulating polycistronic ORs to efficiently induce large-scale olfactory modulation. Indeed, all of the AgORs identified in polycistronic cluster 1B (Karner et al., 2015) are present in our sequencing libraries, and one of these AgORs17 was significantly over-represented in malarial mosquitoes; the increased abundance of the remaining five AgORs (13, 15, 16, 47, 55) are just below statistical significance (Figure 5A).

### Odorant Binding Proteins/Chemosensory proteins

Odorant binding proteins (OBPs), chemosensory proteins (CSPs) and sensory appendage proteins (SAPs) have been postulated to mediate the solubility and transport of odorant molecules within the sensilla lymph (Leal, 2013). A total of 57 AgOBPs/AgCSPs/AgSAPs were identified in sequencing datasets, with 47 AgOBPs/AgCSPs more abundant in heads of infected mosquitoes (17 significantly) while two AgOBPs (9 and 25) and a single AgCSP5 were significantly under-represented relative to uninfected controls (Figure 5B). These findings are consistent with microarray-based transcriptome studies in *Ae. aegypti* that determined dengue viral infection to increase abundance of a subset of OBPs that was correlated to increased host feeding and vectorial capacity (Sim et al., 2012). It is reasonable to postulate that *Plasmodium* infection similarly impacts anopheline mosquitoes to potentially increase host feeding, synergistically supporting parasite and mosquito survivorship as well as maximizing vectorial capacity. AgCSP5 (-0.72-fold; p-adj = 0.014; AGAP029127), AgOBP9 (-0.54-fold; p-adj = 0.030; AGAP000278), and AgOBP25 (-0.40-fold; p-adj = 0.018; AGAP012320) are the only significantly under-represented (-1.77-fold; ACOM030596) OBP/CSP transcripts (Figure 5B). In any case, the precise role of AcCSP5, AgOBP9, and AgOBP25 along with those of other OBP/CSP genes in *An. gambiae* olfaction remains enigmatic.

### Gustatory Receptors

Phylogenetic-based annotation (Kent et al. 2008; Sparks et al. 2013) along with appendage-specific transcriptome profiling of *An. coluzzii, An. gambiae,* and *Ae. aegypti* gustatory receptors (GRs) correlate several orthologous GR genes with the gustatory responses of putative gustatory receptor neurons (GRNs) on the proboscis and labellum. That said, there is a paucity of direct functional data for the majority of *An. gambiae* gustatory receptors (AgGRs) (Matthews et al., 2016; Pitts et al., 2011; Rinker et al., 2013a). In contrast, AgGR22 (AGAP00999), AgGR23 (AGAP003098), and AgGR24 (AGAP001915), which make up a complex that is required for volatile carbon dioxide (CO2) detection critical for blood-feeding behaviors, have been well characterized (Liu et al., 2020). The differential transcriptome profiles for AgGR22 (0.28-fold), AgGR23 (0.31-fold), and AgGR24 (0.34-fold) are only slightly more abundant in sporozoite-infected head transcriptomes than in controls (Figure 7A, red bar) and, not surprisingly, these changes fail to meet the rigorous significance thresholds employed here. This suggests that *Plasmodium* infection does not significantly impact CO2 sensitivity and is consistent with previous electrophysiology-based studies that showed *P. berghei* infections had no effect on *An. stephensi* maxillary palp neuron sensitivity to CO2 (Grant et al., 2013). Taken together, it appears likely that *P. falciparum* parasites do not manipulate early, long-range anopheline host-seeking behaviors, such as CO2 detection (Nguyen et al., 2017).

**Figure 7.**
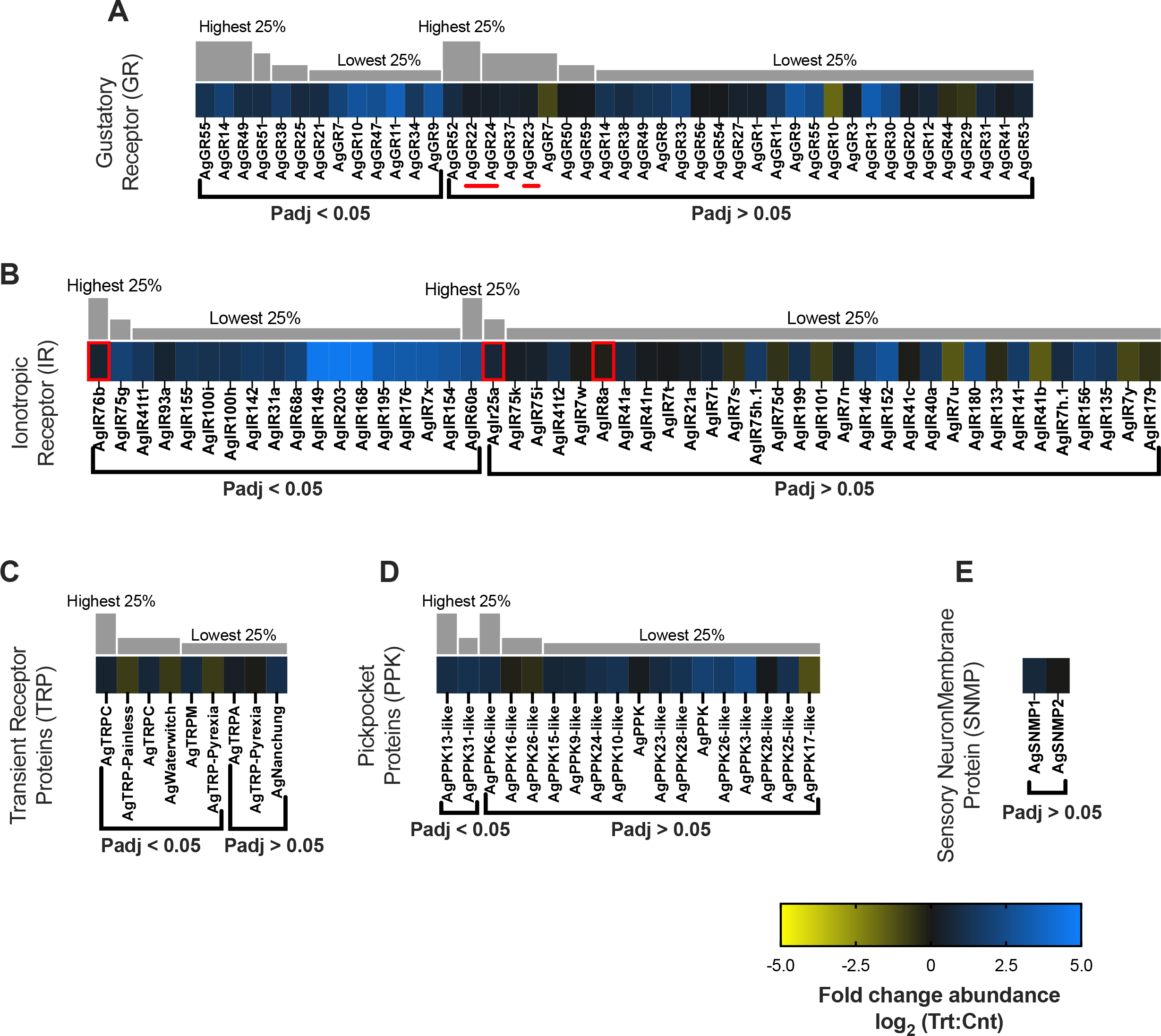
Chemosensory differential transcript abundances following relevant sporozoite infections. Chemosensory transcripts that were present at significantly higher (blue) or lower (yellow) levels in *P. falciparum* sporozoite infected treatments; non-differentially expressed chemosensory transcripts are denoted as zeros (black). Chemosensory genes within each family organized by adjusted p-value and subsequently arrayed left to right from most abundant to least abundant based on FPKM values (quartile bars above each image). (A) Gustatory receptor family (GR). (B) Ionotropic receptor family (IR). (C) Transient receptor protein family (Trpis et al.). (D) Pickpocket protein family (Thiery et al.). (E) Sensory neuron membrane protein family (SNMP). Chemosensory co-receptors are identified by red boxes. The three CO2 gustatory receptors are identified by a red line. Log2 scale indicates transcript abundances that were significantly higher (blue) or lower (yellow) in sporozoite infected treatments (Vrba *et al*.) vs. naïve controls (Cnt).

Beyond the CO2 receptors, we observed a considerable shift in the AgGR transcriptome profile between heads of uninfected and those of sporozoite-infected mosquitoes. A total of 44 AgGRs were differentially detected; transcripts for 40 AgGRs showed a higher abundance and the remaining 4 were less abundant in heads of sporozoite-infected mosquitoes than in controls (Figure 7A). Of these differentially abundant AgGRs, 13 were present at significantly higher levels and none at significantly lower levels in treatments. While the most significantly abundant, AgGR11 (3.43-fold), and seven other AgGRs that are significantly more abundant in treatments remain unannotated, several AgGRs with homology to functionally characterized sugar-responding GRs from *Drosophila* display significantly higher transcript abundances than controls, suggesting a higher sensitivity to sugary substrates in treatments. These include the DmGR5a orthologs AgGR14 (1.94-fold; p-adj = <0.001; AGAP006399) and AgGR21 (0.88-fold; p-adj = 0.05; AGAP003260); the DmGR43a ortholog AgGR25 (0.83-fold; p-adj = 0.05; AGAP004727); and the DmGR68a ortholog AgGR9 (2.90-fold; p-adj = <0.0001; AGAP009805; Figure 7A) (Kent et al., 2008). It is reasonable to hypothesize that because glucose is the primary energy source for both mosquitoes and the *Plasmodium* sporozoites, it would be advantageous to manipulate its acquisition, uptake and transport, thereby synergistically supporting sporozoite and mosquito health, survival and by extension, malaria transmission (Gary and Foster, 2001; Pumpuni and Beier, 1995). This differential increase in putative sugar-responsive AgGRs as a consequence of salivary gland *P. falciparum* infections also correlated with a significant increase in the abundance of several sugar transporters (Supplementary Infomation). While this is the first identification of a *P. falciparum*-mediated increase in abundance of non-CO2 responsive AgGRs, our data are consistent with previous studies highlighting the overabundance of sugar transporters in the salivary glands of *Plasmodium*-infected *An. gambiae* and their importance for sporozoite survival (Pinheiro-Silva et al., 2015). The remaining transcript, AgGR47 (2.60-fold; p-adj = 0.05; AGAP005514), found significantly abundant in treatments is a DmGR66a ortholog and putatively functions in bitter/caffeine detection (Kent et al., 2008). It is reasonable to speculate that *Plasmodium* sporozoites in *Anopheles* mosquitoes upregulate the deterrent effects of such AgGR neuronal responses to prevent ingestion of detrimental bitter nutrient sources that have been shown to temporally inhibit mosquito sugar and water neurons and reduce nectar feeding of *An. quadrimaculatus* (Sparks and Dickens, 2016).

Shifting mosquito gustatory profiles to increase sensitivity to bitter and behaviorally aversive semiochemicals therefore increases the likelihood of blood-meal (and or nectar) feeding and consequently pathogen transmission. It is likely that this effect contributes to increased sensitivity and the marked decrease in mortality rate of *An. gambiae* harboring *P. falciparum* sporozoites when exposed to DEET (N, N-diéthyl-3-méthylbenzamide), a widely used repellent, in comparison with uninfected *An. gambiae* and other pyrethroid test substrates (Mulatier et al., 2018; Swale et al., 2014). Lastly, the increased sensitivity of AgGR47 observed in sporozoite-infected *An. gambiae* relative to naïve controls may reflect age-related nutrient deficiencies in controls. In aging *Drosophila*, nutrient deprivation dramatically alters feeding behaviors, including the depotentiation of responses to bitter (Maruzs et al., 2019).

### Ionotropic Receptors

Ionotropic receptors are an ancient family of chemosensory receptors evolutionarily derived from ionotropic glutamate receptors (Croset et al., 2010) that are associated with olfactory responses to amines and carboxylic acids that have been identified as important semiochemicals closely associated with host-preference and seeking (AgIRs; Montell and Zwiebel, 2016). As is true in other insects, AgIR8a, AgR25a and AgIR76b likely function as co-receptors along with odorant-recognizing AgIRs that ’tune’ responses to host cues (Liu et al., 2010; Pitts et al., 2017). In *An. gambiae*, AgIR25a together with AgIR76b and AgIR8a function in the detection of amines and carboxylic acids, respectively (Pitts et al., 2017) both of these chemical classes have been directly implicated in anopheline host seeking (Smallegange et al., 2005; Takken and Verhulst, 2013). Interestingly, the abundance of AgIR76b transcripts was significantly enhanced (0.55-fold; p-adj = 0.05; AGAP011968) in infected mosquito heads while the levels of AgIR25a and AgIR8a were not significantly different (Figure 7B). Transcripts for 38 of the 49 odorant-tuning AgIRs detected here were found to be more abundant in sporozoite-infected head transcriptomes; only 11 were downregulated compared with uninfected controls (Figure 7B). Of those upregulated, 18 were significantly more abundant but none were significantly less abundant in treatment groups than controls. Inasmuch as many AgIRs are uncharacterized, including those with the most pronounced and significant differential abundance (AgIR149, AgIR168, and AgIR7203), the implications of these differentially up-/downregulated IRs are difficult to assess. Nevertheless, functional information for a small subset of AgIRs is available and others can be derived from analyses of clear orthologs in *Ae. aegypti, An. gambiae, An. sinensis* and *Drosophila melanogaster*. This allows us to identify several AgIRs significantly more abundant in treatment groups (Figure 7B) to be associated with both gustation [AgIR7x (3.11-fold; p-adj = 0.03; AGAP013520), AgIR93a (0.52-fold; p-adj = 0.05; AGAP000256) (Hill et al., 2019), AgIR60a (2.48-fold; p-adj = 0.05; AGAP011943) (Menuz et al., 2014)] and host-seeking behaviors [AgIR75g (1.96-fold; p-adj = 0.003; AGAP013085) (Hill et al., 2019)]. In addition, we observed three receptors localized to the antenna [AgIR31a (1.43-fold; p-adj = 0.03; AGAP009014), AgIR68a (2.00-fold; p-adj = 0.003; AGAP007951), and AgIR41t1 (1.44-fold; p- adj = 0.03; AGAP004432)(Wang et al., 2018)]. While persistence of host seeking and blood feeding might well be advantageous for *Plasmodium,* it would only be beneficial for their anopheline hosts if these behavioral shifts promoted the acquisition of a successful blood meal.

### Transient Receptor Potential Channels

As is true for all insects, *An. gambiae* transient receptor potential (AgTRP) family members are associated with a variety of sensory modalities such as chemosensation, gustation, thermosensation, hygrosensation, mechanosensation, vision, and intracellular signaling (Montell and Zwiebel, 2016). A total of nine AgTRPs were identified in sequencing libraries, five of which are more abundant (three significantly) in heads of sporozoite-infected mosquitoes while four AgTRPs (three significantly) are under-represented relative to uninfected controls (Figure 7C). Of these, AgTRPC (canonical; 0.50-fold; p-adj = 0.0002; AGAP000349), AgTRPC (canonical; 0.67-fold; p-adj = 0.0004; AGAP008435), and AgTRPM (melastatin; 0.75-fold; p-adj = 0.01; AGAP006825) are all active in thermosensation and feeding. TRPC has been identified as having several physiological roles in insects, including signal transduction, phototransduction, and gustatory sensitivity to CO2 (Badsha et al., 2012). TRPM mediates *Drosophila* nociception behaviors and putatively critical behaviors for *An. gambiae* viability and survival (Fowler and Montell, 2013). Given the critical roles that AgTRPC and AgTRPM are likely to play in mosquito host seeking and blood feeding, manipulation of these transcripts would likely be advantageous for *Plasmodium* to ensure maximum host contact rate and transmission. TRP-Painless and TRP-Pyrexia function in thermosensation and have been documented to mediate avoidance of noxious high temperatures in *D. melanogaster* (Sokabe and Tominaga, 2009). It is possible the significantly reduced abundance of AgTRP-Painless (-0.78-fold; p-adj = 0.01; AGAP001243) and AgTRP-Pyrexia (-0.75-fold; p-adj = 0.0003; AGAP010269) in treatment groups compared with naïve controls is related to a loss of nociception.

The remaining *An. gambiae* TRP identified as significantly differentially abundant between treatments and controls functions as a mechanoreceptor with roles in *Drosophila* hygrosensation. AgTRP- Waterwitch which is significantly less abundant in treatment tissues (-0.70-fold; p-adj = 0.001; AGAP000361) is homologous to the DmTRP-Waterwitch that functions in dry-air detection (Bohbot et al., 2014). Moisture detection, or importantly the lack thereof, mediates flight and host-seeking behavior in anopheline mosquitoes such that augmenting mosquito traps with moist air significantly increases flight activity and attraction of wild *Aedes, Anopheles* and *Culex* mosquitoes (Cribellier et al., 2020). It is likely that upregulation of chemo/thermosensory TRPs as well as the inverse regulation of Ag-Waterwitch optimizes persistent host seeking and blood feeding and is consistent with studies showing that *Anopheles* mosquitoes harboring *Plasmodium* sporozoites exhibit more persistent host-seeking behavior than uninfected controls (Ferguson and Read, 2004).

### Pickpocket Channel Proteins/Sensory Neuron Membrane Proteins

We identified 18 *An. Gambiae* pickpocket channel proteins (AgPPKs) of which transcripts were predominately more abundant in heads of sporozoite-infected mosquitoes (Figure 7D). Two of these AgPPKs, AgPPK13-like (0.97-fold; p-adj = 0.004; AGAP007945) and AgPPK31-like (1.47-fold; p-adj = 0.05; AGAP000657), showed significant enrichment. In *D. melanogaster*, PPKs have been implicated to function in salt and water taste, although the behavioral implications of these characterized PPKs is unclear (Freeman and Dahanukar, 2015). Similarly, while aedine PPKs are hypothesized to function in a variety of sensory modalities, such as chemosensation, hygrosensation and mechanosensation (Matthews et al., 2016), anopheline PPKs remain largely uncharacterized and the implications of these differentially abundant PPKs remain unclear. Sensory neuron membrane proteins (SNMPs) are both colocalized with ORs and broadly expressed throughout *D. melanogaster* and are postulated to have both chemosensory and non-chemosensory functions (Vogt et al., 2021) with particular roles in dipteran and lepidopteran pheromone detection (Cassau and Krieger, 2020). Both AgSNMP1 and AgSNMP were identified in sequencing libraries, although without significance (Figure 7E). These and other anopheline SNMPs remain uncharacterized, and the implications of differentially abundant AgSNMPs are therefore unclear.

### General Neuronal Function

Insect neurotransmitters comprise acetylcholine (ACh), gamma-aminobutyric acid (GABA), glutamate (Glu), and select biogenic amines such as dopamine, histamine, octopamine, serotonin and tyramine as well as their respective membrane receptors (Osborne, 1996). In total, 11 subunits for nicotinic ACh receptors (nAChr) and muscarinic ACh receptor subunits (mAChr) were identified with 5 Ag-nAchr and 1 Ag-mAchr significantly overabundant in the heads of sporozoite-infected *An. gambiae*: Ag-nAChr2 (0.75-fold; p-adj = 0.001; AGAP004675), Ag-nAChr⍺2 (0.96-fold; p-adj = <0.0001; AGAP002972), Ag-nAChr⍺3 (1.18-fold; p-adj = <0.0001; AGAP00329), Ag-nAChr⍺5 (1.25-fold; p-adj = <0.0001; AGAP008588), Ag-nAChr⍺6 (1.731-fold; p-adj = <0.0001; AGAP002152), and Ag-nAChrβ1 (0.90-fold; p-adj = <0.0001; AGAP000966; Figure 8A). Only Ag-nAChr⍺9 (-1.23-fold; p-adj = <0.0001; AGAP009493) was significantly diminished in heads of sporozoite-infected mosquitoes compared with naïve mosquitoes (Figure 8A). While insect and especially mosquito AChRs (nicotinic and muscarinic) are not well characterized, it has been reported that with age, AChR responses become significantly diminished as a function of oxidative damage and neurodegeneration (Araujo et al., 1990). In *Drosophila* this is associated with aged-related immune system dysfunction (Nainu et al., 2019). As such, the increased abundance of AgAChrs in sporozoite-infected *An. gambiae* suggests that malaria infected anopheline mosquitoes are resistant to both aged-related immune system dysfunction and neurodegeneration. A similar logic could apply to the significant overabundance of acetylcholinesterase (Ace; AgAce, 1.18- fold; p-adj = <0.0001; AGAP000466) identified in treated *An. gambiae* relative to naïve controls (Figure 9A). As such, the increased levels of AChRs in treatment groups could also represent altered baseline levels within mosquitoes that as a result are not yet susceptible to the effects of aging and senescence.

**Figure 8.**
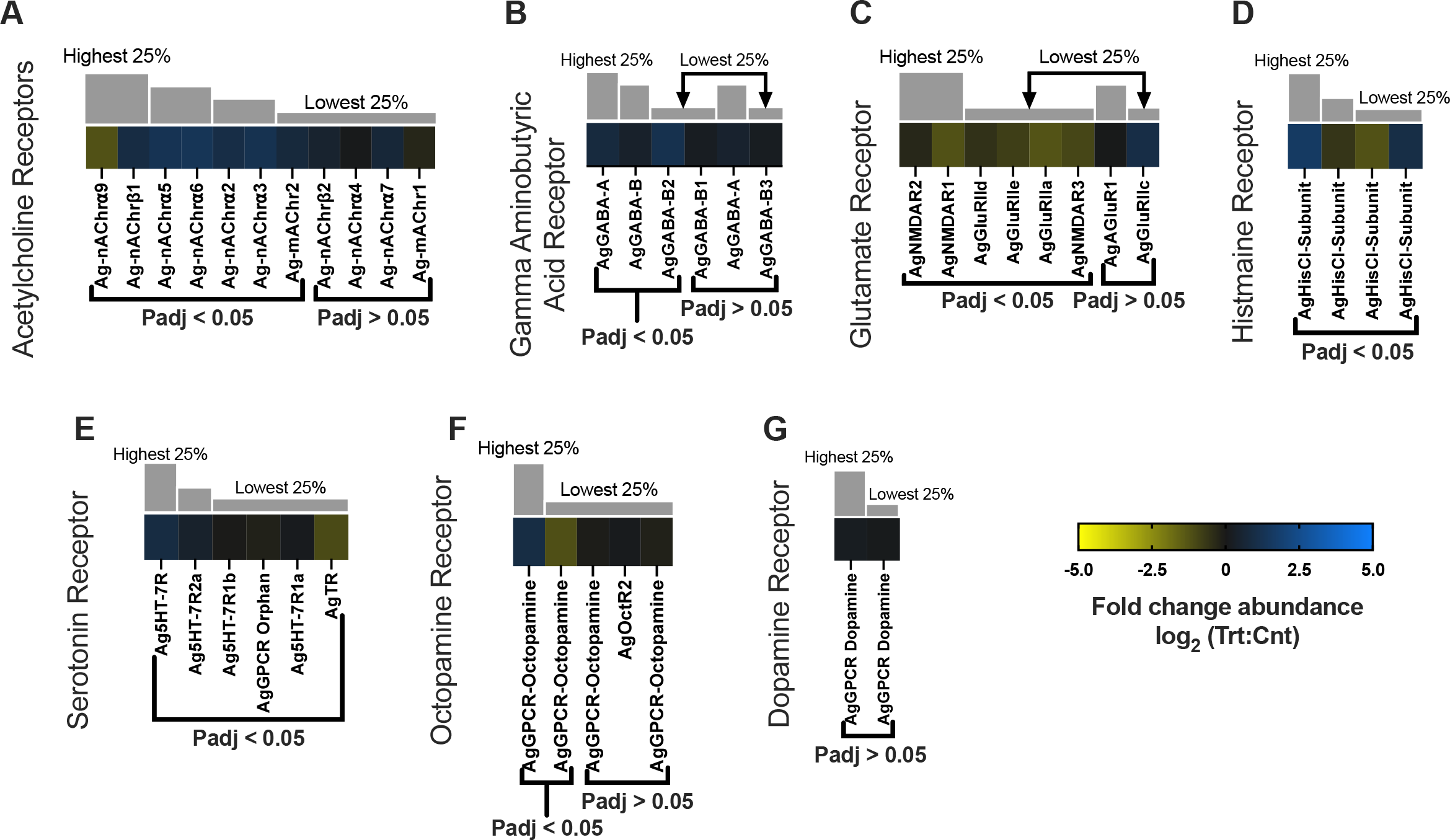
Neurotransmitter differential transcript abundances following relevant sporozoite infections. Neurotransmitter transcripts that were present at significantly higher (blue) or lower (yellow) levels in *P. falciparum* sporozoite infected treatments; non-differentially expressed neurotransmitter transcripts are denoted as zeros (black). Neurotransmitter genes within each family organized by adjusted p- value and subsequently arrayed left to right from most abundant to least abundant based on FPKM values (quartile bars above each image). (A) Acetylcholine receptor family. (B) Gamma aminobutyric acid receptor family. (C) Glutamate receptor family. (D) Histamine receptor family. (E) Serotonin receptor family. (F) Octopamine receptor family. (G) Dopamine receptor family. Log2 scale indicates transcript abundances that were significantly higher (blue) or lower (yellow) in sporozoite infected treatments (Vrba *et al*.) vs. naïve controls (Cnt).

**Figure 9.**
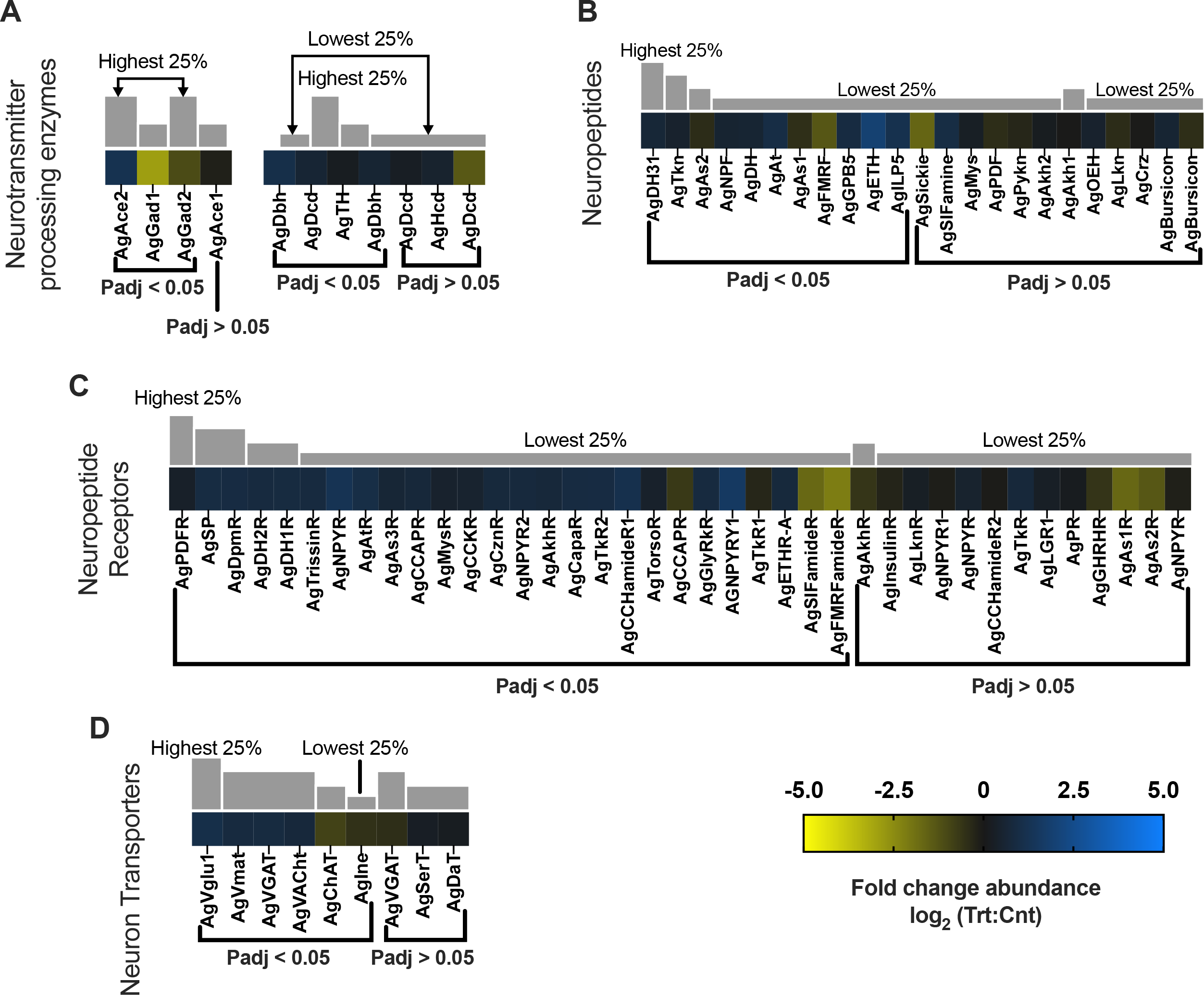
Neuropeptide differential transcript abundances following relevant sporozoite infections. Neuropeptide transcripts that were present at significantly higher (blue) or lower (yellow) levels in *P. falciparum* sporozoite infected treatments; non-differentially expressed neuropeptide transcripts are denoted as zeros (black). Neuropeptide genes within each family organized by adjusted p-value and subsequently arrayed left to right from most abundant to least abundant based on FPKM values (quartile bars above each image). (A) Neurotransmitter processing enzyme family. (B) Neuropeptide family. (C) Neuropeptide receptor family. (D) Neuron transporter family. Log2 scale indicates transcript abundances that were significantly higher (blue) or lower (yellow) in sporozoite infected treatments (Vrba *et al*.) vs. naïve controls (Cnt). ***Finishing Last heat map edits!

Transcripts for both ionotropic (GABA-A) receptors and metabotropic (GABA-B) receptors were significantly more abundant in treatment groups when compared with naïve controls: AgGABA-A (0.80-fold; p-adj = 0.0007; AGAP006028), AgGABA-B (0.38-fold; p-adj = 0.02; AGAP000038), and AgGABA-B2 (1.14-fold; p-adj = <0.0001; AGAP004595; Figure 8B). Transcripts for the GABA processing enzyme glutamate decarboxylase (Gad) displayed a dissimilar differentiation as the abundance of transcripts encoding AgGad1 (-2.92-fold; p-adj = <0.0001; AGAP005866) and AcGad2 (-1.10-fold; p-adj = 0.0001; AGAP008904) were significantly diminished in treatment groups compared with controls (Figure 9A). In *Drosophila*, GABA mechanisms help stabilize neuronal function and maintain tissue homeostasis (Maruzs et al., 2019). As such, the decreased abundance of AgGABAs and increased abundance of AgGads in naïve controls may actually reduce aging-related neurodegeneration to help maintain neuronal homeostasis. Similarly, transcripts for NMDA and non-NMDA types of anopheline glutamate receptors were identified in sequencing libraries and predominately under-represented in heads of infected *An. gambiae*. Of the eight glutamate receptor transcripts identified, none was significantly abundant in heads of sporozoite-infected mosquitoes; in contrast, six NMDA and non-NMDA glutamate receptors—AgGluRIIa (-1.27-fold; p-adj = <0.0001; AGAP000803), AgGluRIId (-0.54-fold; p-adj = 0.003; AGAP002797) AgGluRIIe (-0.81-fold; p-adj = <0.0001; AGAP012447), AgNMDAR1 (-1.23-fold; p-adj = <0.0001; AGAP001478), AgNMDAR2 (-0.32-fold; p-adj = 0.005; AGAP012429) and AgNMDAR3 (-0.98-fold; p-adj = 0.002; AGAP005527)—were significantly diminished in sporozoite-infected mosquitoes compared with naïve controls (Figure 8C). It is possible that in light of the ability of these receptors to function as immunomodulators of inflammatory responses (Hodo et al., 2020), the overabundance of NMDA and non-NMDA glutamate receptors in naïve mosquitoes reflects either an aging-related immune system dysfunction or neuronal pathology. Transcripts encoding the receptors for several peptide biogenic amines, including dopamine, histamine, octopamine, serotonin and tyramine, which function as anopheline neuromodulators and neurotransmitters were identified in our libraries. With the exception of two pairs of histamine and octopamine receptor transcripts that were similarly up-/downregulated, most biogenic amine receptors showed no significant differential abundance in heads of sporozoite-infected *An. gambiae* relative to naïve controls (Figure 8D-G).

Transcripts encoding several biogenic amine processors were identified in similarly up- /downregulated pairs with no clear distinction between treatment and naïve controls (Figure 9A). Insect neuropeptides are essential neuromodulators that regulate a variety of physiological and behavioral functions. Several neuropeptide transcripts identified in sequencing libraries were shown to be significantly overabundant in the heads of sporozoite-infected *An. gambiae* (Figure 9B). Interestingly, there were also four neuropeptide transcripts and their correlating receptors that were significantly diminished in treatment heads compared with naïve controls: allatostatin 1 (AgAs1; -0.50- fold; p-adj = 0.001; AGAP003712), allatostatin 2 (AgAs2; -0.41-fold; p-adj = 0.01; AGAP010157), FMRFamide (AgFMRF, -1.34-fold; p-adj = <0.0001; AGAP005518) and sickie (AgSickie; -1.66-fold; p- adj = <0.0001; AGAP009424) were directly or indirectly correlated with insect stress and immune response (Figure 9B-C). In *Ae. Aegypti*, allatostatin interacts with hemocytes to increase humoral immune responses to combat cellular stress and inflammation (Hernandez-Martinez et al., 2017). FMRFamide signaling in *Drosophila* increases in response to cellular stress and inflammation to promote stress-induced sleep and cellular recovery to help bolster immune function (Lenz et al., 2015). Sickie acts directly on *Drosophila* innate immunity by activating the transcription factor Relish and subsequent transcription of immune defense antimicrobial peptides (AMPs; Govind, 2004 #31754). Neuropeptide F (NPF) also acts indirectly to modulate stress responses through activation of downstream signaling pathways. In dipterans, NPF independently activates insulin signaling pathways and coordinately the target or rapamycin (TOR) pathway (Dhara et al., 2013). TOR pathway is an insect nutrient-sensing pathway that regulates cellular metabolism and homeostasis in response to nutrient flux and stress (Lee et al., 2008; Wullschleger et al., 2006). TOR signaling naturally increases with age and increasing cellular dysfunction and stress (Katewa and Kapahi, 2010). With a substantial number of neuropeptides/hormones and their correlating receptors significantly abundant in heads of sporozoite-infected *An. gambiae* and functioning to maintain healthy mosquito cells and tissues, the decreased abundance of these select five neuropeptides in infected mosquitoes is puzzling. If the “normal’ transcriptome profile of naïve mosquitoes reflect changes associated with aging-related immune system dysfunction and the loss of cellular homeostasis, or some other age-related pathology, then one could hypothesize that infected mosquitoes might be refractory to those effects.

### Immune System

Activation of the immune system in response to infection is energetically costly and likely results in decreased fitness reflected by other life history traits with the potential to impact vectorial capacity. Mosquito anti-*Plasmodium* immunity is stage specific and involves diverse defense processes, although the extent of immune defense in salivary-gland-stage infected mosquitoes, and especially the head and salivary glands (post sporozoite salivary gland invasion), remains unclear (Belachew, 2018). Since *Plasmodium*, unlike arboviruses, does not establish infection in the mosquito head, the immunity-related transcriptome changes observed in our study are likely to mostly derive from the combined head (including sensory appendages) and salivary gland transcriptomes. In heads of sporozoite-infected and control sequencing libraries, several components of *An. gambiae* humoral and cellular, innate immune defenses were identified. *An. gambiae* Pattern Recognition Receptors (PRRs) include 6 fibrinogen proteins (FBGs), 7 3-glucan binding proteins (3GNBPs), 7 peptidoglycan recognition proteins (PGRPs), 24 leucine-rich repeat immune proteins (LRIMs) including *Plasmodium* infection responsive leucine-rich proteins, 13 thioester-containing proteins (TEPs), and 2 members of C-type lectin family proteins (CTLs)—AgCTL4 and AgCTLM2 (Figure 10A, 10B). A large portion of the PRR transcripts were not present at significantly differential abundances between heads of sporozoite-infected and control mosquitoes; indeed, neither of the transcripts encoding AgCTLMA2 or AgCTL4 displayed significant differential abundance.

**Figure 10.**
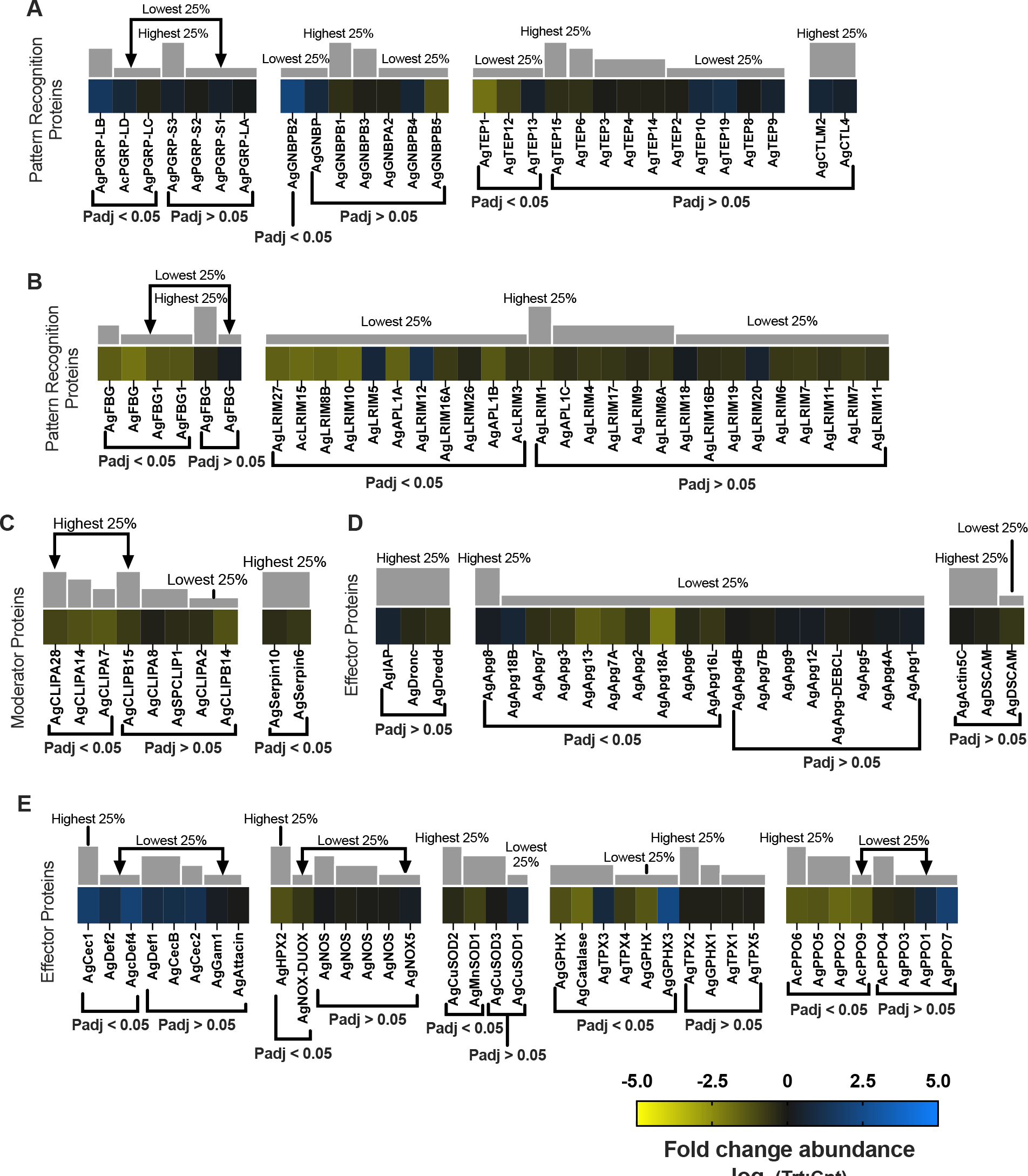
Immune response differential transcript abundances following relevant sporozoite infections. Immune response transcripts that were present at significantly higher (blue) or lower (yellow) levels in *P. falciparum* sporozoite infected treatments; non-differentially expressed immune response transcripts are denoted as zeros (black). Immune response genes within each group organized by adjusted p-value and subsequently arrayed left to right from most abundant to least abundant based on FPKM values (quartile bars above each image). (A-B) Pattern recognition proteins. (C) Modulator proteins. (D-E) Effector proteins. Log2 scale indicates transcript abundances that were significantly higher (blue) or lower (yellow) in sporozoite infected treatments (Vrba *et al*.) vs. naïve controls (Cnt).

Anopheline TEPs are antimicrobial and antiparasitic PRRs, complement-like factors that drive pathogen clearance through lysis and melanization (Blandin et al., 2004). Of those, only AgTEP13 (0.34-fold; p-adj = 0.03; AGAP008407) was significantly more abundant in heads of sporozoite-infected mosquitoes, and AgTEP1 (-1.95; p-adj = <0.0001; AGAP010815) and AgTEP12 (-0.94; p-adj = 0.011; AGAP008654) significantly diminished in treatment groups compared with controls (Figure 10A). While the roles of TEP12 and TEP13 in anopheline immune responses are not currently known, the well-studied TEP1 functions as an enhancer of mosquito phagocytosis (Kumar et al., 2018). Since TEP1 is incapable of colocalizing and killing *Plasmodium* sporozoites, its decreased abundance in treatment groups is not unexpected (Blandin et al., 2004) (Yan and Hillyer, 2019). As such, the increased abundance of AgTEP1 in naïve controls likely reflects aging-related immune system dysfunction, as aging decreases phagocytic efficiency and contributes to immunosenescence (Maruzs et al., 2019). LRIMs and APLs are PPRs that mediate innate immune surveillance and response. Transcripts for 20 LRIMs and APLs were downregulated in heads of infected mosquitoes relative to naïve controls, while 4 of the 24 *An. gambiae* LRIMs were more abundant in treatments. Of the 4 AgLRIMs upregulated, 2 were significantly more abundant, whereas 9 of the 20 LRIMs and APLs were significantly less abundant in treatments than controls (Figure 10B). These data are largely consistent with previous studies on the transcriptional regulation of LRIMs and APLs in response to *Plasmodium,* gram-negative/gram-positive bacteria, and fungal challenge (Aguilar et al., 2005; Waterhouse et al., 2010). While AgLRIM12 (0.72-fold) and AgLRIM5 (0.82-fold; ACOM043147) were both significantly more abundant in heads of infected mosquitoes they are not yet conclusively associated with *Plasmodium* infection (Dong et al., 2006a; Waterhouse et al., 2010); it is therefore possible the observed differential abundance of AgLRIMs reflect non-immunity responses or potential aging effects that are only partially impacted by *P. falciparum* infection.

Similarly identified AgFBGs, which function in phagocytosis and complement activation, were predominately diminished in treatments relative to naïve controls, with AgFBG (-1.47-fold; p-adj = 0.015; AGAP011231), AgFBG (-1.98-fold; p-adj = 0.004; AGAP010869), AgFBG (-1.29-fold; p-adj = 0.02; AGAP004917) and AgFBG (-1.29-fold; p-adj = 0.007; AGAP006914) significantly reduced (Figure 10B; Kumar et al., 2018). In *Drosophila,* aging is associated with increased microbial loads and dysbiosis that trigger increased expression of immune genes and stronger immune activation (Garschall and Flatt, 2018). Excessive immune activation ultimately results in immune dysfunction and decreased lifespan (Guo et al., 2014). In that context, the large number of AgLRIMs, AgAPLs and AgFBGs that are more abundant in naïve control transcriptomes may reflect aging-related dysfunction of the *An. gambiae* immune system. Accelerated immune system dysfunction in naïve controls also rationalizes the observed AgPGRP transcript abundances identified in replicate sequencing libraries. Our data indicate that transcripts for two AgPGRPs that have been characterized as negative regulators of immune IMD signaling defense (Ramirez et al., 2020)—AgPGRP-LB (1.46-fold; p-adj = 0.02; AGAP001212) and AgPGRP-LD (0.69-fold; p-adj = 0.05; AGAP005552: Figure 10A)—are significantly more abundant in heads of sporozoite-infected mosquitoes, suggesting a suppression of the IMD immune pathway. The IMD signaling initiation receptor AgPGRP-LC (-0.25-fold; p-adj = 0.02; AGAP005203), identified as significantly reduced in treatments compared with controls, further supports the presence of diminished IMD signaling in sporozoite-infected *An. gambiae*. PGRP immune regulator deficiency results in unregulated IMD signaling causing dysplasia, inflammation, apoptosis, and decreased lifespan (Gupta et al., 2009). Preservation of immune homeostasis would be advantageous for both *Anopheles* hosts and *Plasmodium* pathogens to maintain vectorial capacity. It is possible that *An. gambiae* challenged with biologically relevant, low-level *P. falciparum* results in a mild and prolonged engagement of the mosquito immune system that ultimately primes the system for protection from secondary infection and importantly, the negative impact of aging-related immune system degeneration (Cooper and Eleftherianos, 2017). Indeed, *Plasmodium*- induced anopheline immunological priming may represent an advantageous selective adaptive pressure that maximizes mosquito host fitness and, consequently, vectorial capacity.

Serine proteases constitute the majority of mosquito immune system moderators that are activated in response to pathogen recognition (Kumar et al., 2018). While there are hundreds of serine proteases functioning in a variety of mosquito molecular processes, eight CLIP-domain serine proteases (CLIPs)—SPCLIP1, CLIPA2, CLIPA7, CLIPA8, CLIPA14, CLIPA28, CLIPB14, and CLIPB15— mediate *P. berghei* melanization (Nakhleh et al., 2017; Kumar *et al*., 2018; Sousa et al., 2020). No AgCLIP transcripts were significantly more abundant in heads of sporozoite-infected mosquitoes, although eight were less abundant in treatment groups for which mRNAs of AgCLIPA7 (-1.34-fold: p- adj = <0.0001; AGAP011792), AgCLIPA14 (-1.2-fold; p-adj = <0.0001; AGAP011788) and AgCLIPA28 (-0.92-fold: p-adj = 0.02; AGAP010968) were significantly under-represented relative to uninfected controls (Figure 10C). These data were expected because anopheline CLIPs predominately moderate melanization of rodent *Plasmodium* ookinetes and are not associated with immune defenses targeting sporozoites or human *Plasmodium* (Nakhleh et al., 2017). Indeed, the significant abundance of AgCLIPs in uninfected middle-aged *An. gambiae* may result from aging-related immune dysfunction. The absence of elevated levels of AgCLIPs highlights another putative benefit of *Plasmodium-*mediated immune priming. Another way to moderate immune effector mechanisms such as melanization is performed by Serpins (serine protease inhibitors). Levels of two Serpins, AgSerpin6 (-0.64-fold; p-adj = <0.0001; AGAP009212) and AgSerpin10 (-0.42; p-adj = 0.03; AGAP005246), that are upregulated in response to *Plasmodium* ookinetes (Danielli et al., 2003; Abraham et al., 2005) were significantly under-represented compared with uninfected controls (Figure 10C). As melanization is not expected to be active in *An. gambiae* carrying salivary-gland-stage *P. falciparum* infections, there would be no requirement for Serpin moderation, and such genes as Serpin6 and Serpin10 would no longer be required at this timepoint (Pinto et al., 2007). Likewise, putative immune dysfunction in uninfected controls may impair proper regulation of melanization and, as such, Serpin transcription.

A wide range of immune effectors are downstream of PRR-controlled pathways and other recognition pathways that include anti-microbial peptides (AMPs), reactive oxygen species (ROS), reactive nitrogen species (RNS), prophenoloxidase (PPOs), and apoptosis- and phagocytosis-related genes (Kumar et al., 2018). In *Anopheles*, five classes of AMPs including attacins, defensins, diptericins, cecropins, and gambicins have been described (Christophides et al., 2002). In our data set, three defensins (Def), three cecropins (Cec), one gambicin (Gam) and one attacin (Att) are present (Figure 10E). Of these, only two defensins and Cec1 were significantly differentially abundant between heads of sporozoite-infected and uninfected control mosquitoes, with AgDef2 (0.96-fold; p-adj = <0.0001; AGAP004632), AgDef4 (1.96-fold; p-adj = 0.005; AGAP005416) and AgCec1 (1.78-fold; p-adj = 0.001; AGAP000693) are all significantly abundant in treatment groups compared with controls. Interestingly, these shifts plus the significantly increased abundance of AgGNBPB2 (1.46-fold; p-adj = 0.02; AGAP001212; Figure 10A), a PRR that co-localizes *P. falciparum,* are the only other indication of an *An. gambiae* immune response to *P. falciparum* infection in this study. Of these, only AgCec1 is associated with anti-sporozoite activity. AgDef1 and AgDef4 demonstrate antibacterial activity but have not shown any anti-*Plasmodium* activity (Kokoza et al., 2010). Interestingly, studies in *Ae. aegypti* have determined that differential Def abundances can be observed without immune challenge, and often after pathogen clearance, suggesting alternative functions for the AMPs (Bartholomay et al., 2004). As such, the role of AgDefs in response to sporozoite infections remains unclear. ROS and RNS are vital effectors of Anopheline immune defenses active against malaria parasites (Kumar et al., 2018; Molina-Cruz et al., 2008; Oliveira et al., 2012). ROS formation in dipterans is postulated to involve NADPH oxidase (NOX)-based production of superoxide anions that are transformed into H2O2 by superoxide dismutase (SOD) and subsequent production of ROS (Molina-Cruz et al., 2008). NOX is also likely required for heme peroxidase (HPX) and nitric oxide synthase (NOS) activation, both of which potentiate nitration and generations of RNS (Oliveira et al., 2012). Interestingly, the transcripts for all the enzymes associated with ROS/RNS system and their potentiators were largely under-represented in sporozoite-infected head transcriptomes compared with uninfected controls: AgMnSOD1 (-0.88-fold; p-adj = 0.001; AGAP010517), AgCuSOD2 (-0.38- fold; p-adj = 0.006; AGAP005234), AgHPX2 (-1.20-fold; p-adj = <0.0001; AGAP009033) and AgNOX- DUOX (-0.63-fold; p-adj = 0.03; AGAP009978; Figure 10E). In that light, it is logical to expect the significant under-representation of transcripts encoding catalase, glutathione peroxidase and thioredoxin peroxidase detoxifying enzymes observed in heads of sporozoite-infected *An. gambiae* when compared with uninfected controls (Figure 10E).

In *Anopheles* and other mosquitoes, CLIPs activate PPOs, the primary effectors mediating melanization (Lu et al., 2014). In that light, the lack of differentially abundant AgCLIP transcripts aligns with the observation that AgPPOs were predominately under-represented in heads of sporozoite-infected mosquitoes compared with uninfected controls. AgPPO2 (-1.70-fold; p-adj = 0.001; AGAP006258), AgPPO5 (-1.30-fold; p-adj = <0.0001; AGAM012616), AgPPO6 (-1.38-fold; p- adj = 0.002; AGAP004977), and AgPPO9 (-1.50-fold; p-adj = 0.006; AGAP004978) all significantly diminished in *An. gambiae* carrying salivary-gland-stage sporozoites (Figure 10). The higher PPO levels in heads of uninfected mosquitoes is surprising and may reflect a higher background of metabolic stress. However, this pattern is consistent with the fact that melanized sporozoite-stage *Plasmodium* has never been observed. In *Ae. aegypti,* allatostatin neuropeptide interacts with hemocytes and may have subsequently increased AgPPO expression to address dysbiosis and inflammation resulting from IMD destabilization. This finding is particularly interesting because it correlates mosquito immune system fitness with neuropeptide production and has significantly broader implications for aging-related immune system dysfunction. It also provides a plethora of anopheline molecular mechanisms that putatively synergize with *Plasmodium* immune priming, thereby providing a significant advantage that ultimately facilitates malaria transmission.

Apoptosis, autophagy and phagocytosis genes are important effectors in response to viral and pathogen invasion through programmed cell death of infected cells and clearance of cellular debris and pathogen particles (Kumar et al., 2018). In dipterans, apoptosis is induced through the activation of the long caspases, Dronc and Dredd, and inhibition of inhibitors of apoptosis (IAPs) that in turn regulate autophagy genes, with expression of both increasing concurrently (Eng et al., 2016). Transcripts for AgIAP, AgDronc, AgDredd, 18 autophagy-related proteins (AgApgs), and 2 phagocytosis signaling proteins were identified (Figure 10D). AgDronc, AgDredd, and AgIAP were all under-represented in heads of sporozoite-infected mosquitoes when compared with controls, although none significantly (Figure 10D). AgApgs were predominately under-represented in treatment groups compared with controls, with only AgApg8 (0.20-fold; p-adj = 0.03; AGAP002685) and AgApg18B (0.59-fold; p-adj = 0.003; AGAP005910) significantly abundant in treatment groups, and 8 AgApgs significantly diminished in treatment groups (Figure 10D): AgApg2 (-0.57-fold; p-adj = 0.004; AGAP004092), AgApg3 (-0.69-fold; p-adj = <0.0001; AGAP011582), AgApg6 (-0.37-fold; p-adj = 0.02; AGAP003858), AgApg7 (-0.49-fold; p-adj = 0.001; AGAP010303), AgApg7A (-1.14-fold; p-adj = <0.0001; AGAP008637), AgApg13 (-1.48-fold; p-adj = <0.0001; AGAP005715), AgApg16L (-0.67- fold; p-adj = 0.007; AGAP002315), and AgApg18A (-2.10-fold; p-adj = <0.0001; AGAP007970).

Furthermore, the putative phagocytosis signaling proteins, AgActin-5C and AgDSCAM (down-syndrome cell adhesion molecule), were under-represented in treatment transcriptomes, which is not surprising since both AgActin-5C (Sandiford et al., 2015) and AgDSCAM (Dong et al., 2006b) are linked to anti- *Plasmodium* defense at the midgut-stage infection. Their diminished levels in sporozoite-infected mosquitoes suggest that their active involvement in *Plasmodium* clearance is unlikely. Indeed, the general overabundance of apoptosis, autophagy, and phagocytosis genes in heads of uninfected control mosquitoes likely reflects the tissue specificity of our study, since phagocytosis is mediated by hemocytes and not epithelial cells, and/or a loss of cell homeostasis that accompanies aging-related immune system dysfunction and again highlights another putative benefit of immune priming in *Plasmodium*-infected anopheline vectors.

PRRs activate the Toll pathway through cleavage of Spaetzle proteins (SPZs) to activate Dorsal/NK-kB transcription factor (Rel1) and regulate antimicrobial peptides (Kumar et al., 2018). The transcriptome profile of these components also supports an aging-related immune system deregulation associated with malarial treatments that is importantly not seen in uninfected controls.

Transcripts encoding AgRel1 (-0.44-fold; p-adj = 0.002; AGAM009515) along with one Spaeztle, AgSPZ6 (-0.80-fold; p-adj = 0.002; AGAP005126), are both significantly under-represented in heads of sporozoite-infected mosquitoes (Figure 11A). Inasmuch as the Toll pathway has not been linked to defenses against the human malaria parasite *P. falciparum* (Garver et al., 2009; Dong et al., 2011), it is not surprising that the majority of Toll pathway genes were also significantly under-represented in treatments compared with uninfected controls (Figure 11A). In aging *Drosophila,* Toll signaling becomes increasingly dysfunctional, causing inflammation, systemic organismal defects, and metabolic deficiencies that result in shortened lifespans (Garschall and Flatt, 2018). In that light, decreased activation of Toll in the heads of sporozoite-infected mosquitoes is part of what we propose to be a broadly synergistic *Anopheles*-*Plasmodium* paradigm.

**Figure 11.**
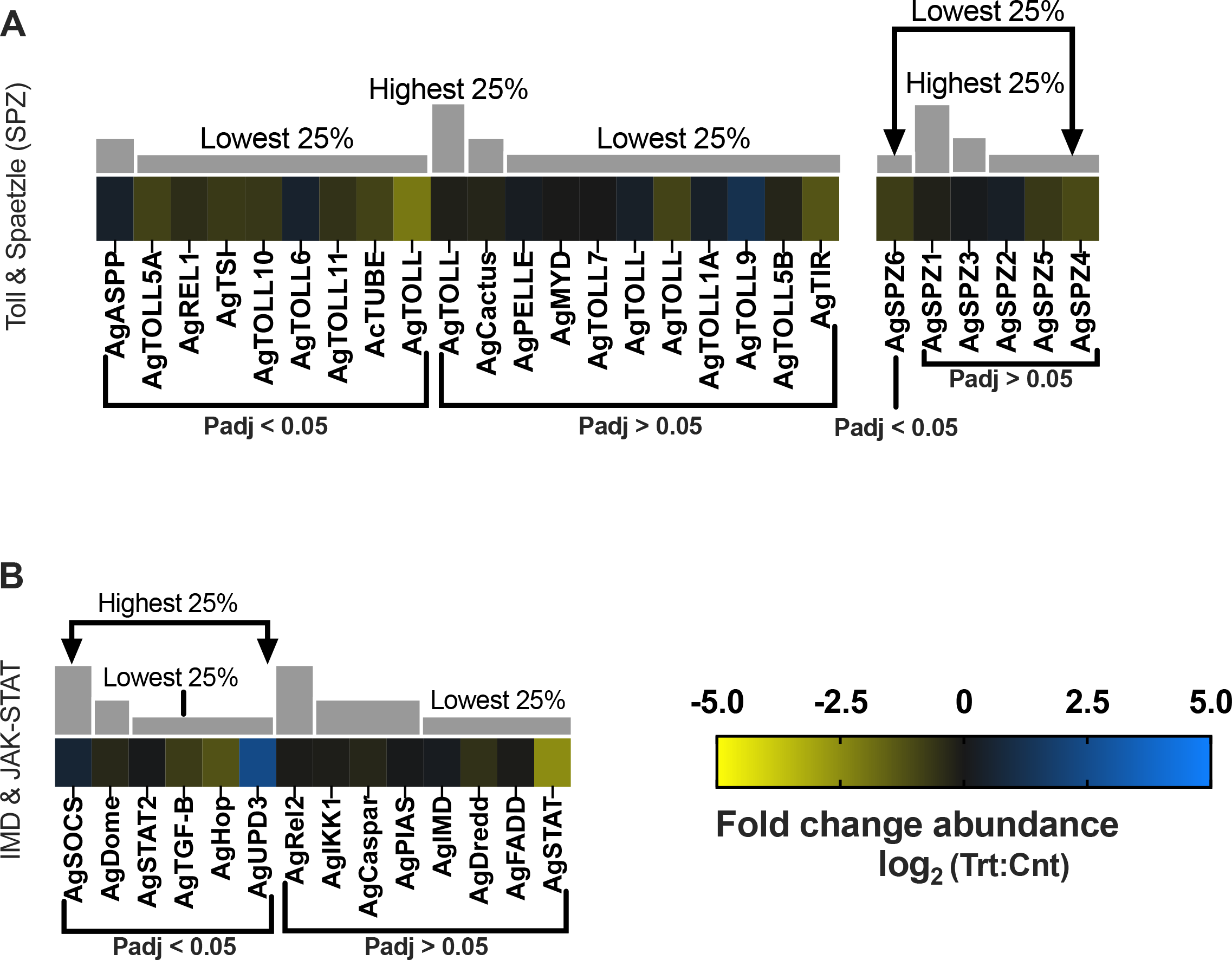
Immune response differential transcript abundances following relevant sporozoite infections. Immune response transcripts that were present at significantly higher (blue) or lower (yellow) levels in *P. falciparum* sporozoite infected treatments; non-differentially expressed immune response transcripts are denoted as zeros (black). Immune response genes within each group organized by adjusted p-value and subsequently arrayed left to right from most abundant to least abundant based on FPKM values (quartile bars above each image). (A) Toll and spaetzle proteins. (B). IMD and JAK-STAT proteins. Log2 scale indicates transcript abundances that were significantly higher (blue) or lower (yellow) in sporozoite infected treatments (Vrba *et al*.) vs. naïve controls (Cnt).

As is the case for Toll-mediated signaling, the anopheline IMD immune signaling pathway involves binding of the PRR, PGRP-LC, which triggers activation of NK-kB transcription factor (Rel2) and production of AMPs. Transcripts for all components of IMD signaling, including PGRP-LC, were identified in sequencing libraries, although AgPGRP-LC (-0.26-fold; p-adj = 0.02; AGAP005203) was the sole transcript significantly diminished in heads of sporozoite-infected mosquitoes (Figure 10A, 11B). Even the modest differential abundance of AgPGRP-LC, which is a positive regulator of IMD signaling and PGRP expression, provides a rationale for the lack of significantly differential IMD signaling pathway transcripts in sporozoite-infected mosquitoes (Figure 11B; Kumar et al., 2018). In any case, the destabilization of anopheline IMD signaling pathway components would putatively render the immune signaling system a poor target for *Plasmodium* immunological priming, as the long-lasting immune response would be unpredictable and difficult to adapt (Cooper and Eleftherianos, 2017).

The anopheline JAK-STAT signaling pathway has been implicated to function in a variety of dipteran developmental processes, as well as in antimicrobial, antiparasitic and antiviral immune defenses (Clayton et al., 2014). It is negatively regulated by PIAS and suppressor of cytokine signaling (SOCS; Shuai and Liu, 2003) and induced by the binding of Unpaired (Upd) to Domeless (Dome) to form a receptor dimer that, in turn, activates Janus Kinase (JAK) or its homolog Hopscotch (Hop) and ultimately STAT which triggers the production of genes regulated by the JAK-STAT pathway and NOS (Kumar et al., 2018). Transcripts for AgSOCS (0.58-fold; p-adj = 0.008; AGAP029624) and AgUPD3 (2.39-fold; p-adj = 0.007; AGAP013506) were significantly more abundant in treatment groups, and AgDome (-0.33-fold; p-adj = 0.006; AGAP029053), AgHop (-1.26-fold; p-adj = 0.003; AGAP008354; JAK homolog) and AgSTAT2 (-0.31-fold; p-adj = 0.02; AGAP000099) were significantly under-represented compared with uninfected controls (Figure 11B). As is the case for IMD pathways, anopheline JAK-STAT signaling primarily targets pre-sporozoite stages of *Plasmodium*, making the lack of JAK-STAT activation in the salivary-gland-stage infected *An. gambiae* examined here expected (Gupta et al., 2009). This is particularly evident by the significant abundance of JAK-STAT’s primary negative regulator AgSOC in heads of sporozoite-infected mosquitoes. Dipteran SOCs deactivate the Hop-Dome dimer, block STAT activation, and target components of the Hop-Dome dimer complex for degradation (Croker et al., 2008). As such, the increased abundance of AgUPD3 in treatment groups may reflect replenishment activity to restore UPD3 levels following *P. falciparum* midgut invasion or simply maintenance of normal UPD3 equilibrium. In *Drosophila*, immune system deregulation and mitochondrial degeneration significantly increase JAK-STAT expression in response to growing systemic inflammation and oxidative stress associated with aging. Excessive JAK-STAT activation ultimately leads to a loss of tissue homeostasis and cell death (Garschall and Flatt, 2018). The active inhibition of JAK-STAT signaling in infected mosquitoes would explain the significantly diminished abundance of AgSTAT2 and AgHop and may also reflect aging-related dysfunction in uninfected *An. gambiae* controls. As such, the anopheline JAK-STAT signaling pathway may represent a target for *Plasmodium* priming, reducing the detrimental effects of aging-related immune deregulation thereby synergistically benefiting both parasite and host.

### General Metabolism and Aging

Metazoan aging can be characterized by the loss of cellular and tissue homeostasis, deterioration of vital biological processes with negative impacts on signaling pathways that regulate nutrient sensing, metabolic allocation, and the maintenance of healthy cells and tissues that together promote mortality (Maruzs et al., 2019). The insect nutrient-sensing pathway TOR regulates cellular metabolism and growth in response to nutrient flux (Wullschleger et al., 2006). In *Drosophila,* increased TOR signaling is associated with aging, and the inhibition of TOR significantly extends fly lifespan and overall viability (Bjedov et al., 2010). Both anopheline TOR transcripts—AgTOR (-0.44-fold; p-adj = <0.0001; AGAP007873) and AgTORC1 (-0.51-fold; p-adj = 0.003; AGAP010035)—are significantly diminished in heads of sporozoite-infected *An. gambiae* compared with naïve controls (Figure 12A). Additionally, a transcript for the TOR kinase inhibitor tuberous sclerosis complex 2 (AgTSC2; -0.46-fold; p-adj = <0.001; AGAP003445), which in *Drosophila* inhibits TOR signaling and acts on fat bodies as a mechanism of lifespan expansion, was found to be significantly diminished in treatment groups compared with controls (Figure 12A; Bjedov *et al*., 2010).

**Figure 12.**
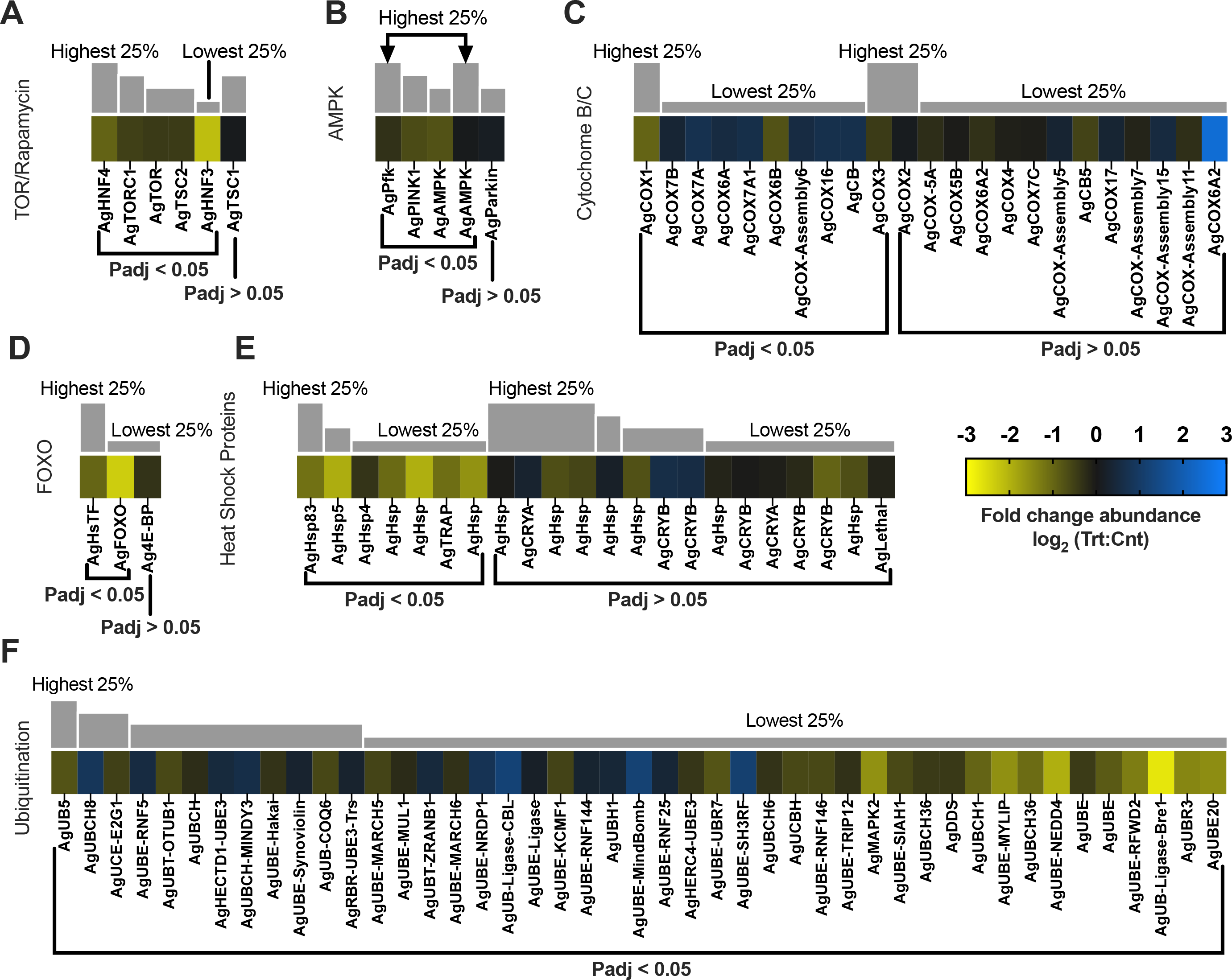
Aging biomarker differential transcript abundances following relevant sporozoite infections. Aging biomarker transcripts that were present at significantly higher (blue) or lower (yellow) levels in *P. falciparum* sporozoite infected treatments; non-differentially expressed Aging biomarker transcripts are denoted as zeros (black). Aging biomarker genes within each group organized by adjusted p- value and subsequently arrayed left to right from most abundant to least abundant based on FPKM values (quartile bars above each image). (A) TOR and rapamycin proteins. (B). AMPK proteins. (C) Cytochrome C and B proteins. (D) FOXO proteins. (E) Heat shock proteins. (F) Ubiquitination proteins. Log2 scale indicates transcript abundances that were significantly higher (blue) or lower (yellow) in sporozoite infected treatments (Vrba *et al*.) vs. naïve controls (Cnt).

The AMP-activated kinase (AMPK) signaling pathway maintains insect energy homeostasis by regulating ATP consumption and production; it is activated in response to low intracellular AMP levels, prompting ATP re-accumulation by reducing costly anaerobic processes as well as activating phosphofructokinase (Pfk), thereby increasing glycogenolysis (Hardie et al., 2012). Interestingly, the transcripts for both AgAMPK (-0.75-fold; p-adj = <0.0001; AGAP002686) and AgPfk (-0.33-fold; p-adj = 0.05; AGAP007642) were significantly diminished in heads of sporozoite-infected *An. gambiae* compared with non-infected mosquitoes (Figure 12B). In contrast, AMPK overabundance in *An. stephensi* is associated with increased innate immune responses and significant alterations in mitochondrial biogenesis (Oringanje et al., 2021). In aging *D. melanogaster*, increased levels of AMPK improve tissue homeostasis via an anti-inflammatory mechanism to promote longevity (Ulgherait et al., 2014). It is possible that, compared with treatment transcriptomes, the age-related increased immune response observed in heads of naïve *An. gambiae* results in reduced nutrient and energy homeostasis. This hypothesis is also supported by the significantly diminished abundance of hepatocyte nuclear factor 3 (AgHNF3; -2.17-fold; p-adj = <0.0001; AGAP001671) and hepatocyte nuclear factor 4 (AgHNF4; -0.98-fold; p-adj = 0.005; AGAP002155) in heads of sporozoite-infected *An. gambiae* mosquitoes compared with naïve controls. In *Ae. Aegypti*, RNAi depletion of HNF4 was shown to result in significant downregulation of transcripts encoding triacylglycerol catabolism, which is essential for fat body metabolism (Wang et al., 2017).

The head transcriptomes of naïve *An. gambiae* controls provide substantial evidence of a robust endemic sterile inflammatory response as well as decreased mitochondrial function, increased mitochondrial damage and mitophagy. In *D. melanogaster,* mitochondrial dysfunction is closely associated with decreased mitochondrial transcript levels of genes such as cytochrome oxidase C (COX) and cytochrome B1 (CB; Schwarze et al., 1998). Indeed, in heads of sporozoite-infected *An. gambiae*, transcripts for COX and CB were present at very high levels, significantly more abundant than in uninfected controls (Figure 12C). In general, naïve *An. gambiae* displayed a significant overabundance of transcripts associated with ubiquitination and autophagy, indicating a higher degree of cell dysfunction than in the treatment group (Figure 12F). For example, PTEN-induced kinase 1 (PINK1), a serine/threonine kinase that specifically targets damaged mitochondria for autophagy, is significantly diminished in *Plasmodium-*infected *An. gambiae* compared with non-infectious mosquitoes (-0.66-fold; p-adj = 0.0004; AGAP004315) (Figure 12B), which may be an indication of a higher degree of mitochondrial dysfunction in naïve aged mosquitoes (Luckhart and Riehle, 2020).

In insects, the forkhead box O (FOXO) transcription factor signaling pathway is an important moderator of protein quality control, stress resistance, and lifespan extension pathways (Maruzs et al., 2019). FOXO functions with its transcriptional target translation initiation factor, 4E-binding protein (4E-BP), to moderate proteostasis in *Drosophila* by removing damaged/abnormal proteins (Demontis and Perrimon, 2010). While Ag4E-BP was not significantly different, AgFOXO (-2.36-fold; p-adj = <0.0001; AGAP000662) was significantly diminished in heads of sporozoite-infected *An. Gambiae* compared with naïve controls. Similarly, the heat-shock transcription factor (AgHsTF; -1.00-fold; p-adj = <0.0001; AGAP029908) and numerous subsequently induced heat shock protein transcripts (Hsps) were all significantly diminished in treatment groups compared with naïve control *An. gambiae* (Figure 12E). In dipterans, protein abnormalities resulting from oxidative stress gradually increase with age and are associated with an increase in HsTF and Hsp production to combat injury from abnormal and malformed proteins (Tower, 2010). Lastly, consistent with these transcriptome shifts, our analyses have also uncovered several transcripts involved in ubiquitin-mediated proteosome protein degradation pathways, the abundance of which are significantly diminished in sporozoite infected mosquitoes (Figure 12F). When viewed collectively, it is likely these differentially abundant transcripts are indicative of an accumulation of abnormal proteins with detrimental effects in naïve, uninfected mosquitoes that importantly does not occur in age-matched counterparts that harbor salivary-gland- stage *P. falciparum* sporozoites after low intensity, biologically relevant blood-meal infections.

## Conclusions

The impact of *Plasmodium* infection on the behavior and physiology of anopheline mosquitoes significantly drives vectorial capacity and this study represents the first comprehensive examination of the effects of biologically relevant *Plasmodium* sporozoite-stage infections on anopheline vectors. To gain insight into this process in as biologically relevant a manner as possible, we have carried out a set of comprehensive comparative analyses of the transcriptome profile of the head (including sensory appendages) and salivary gland tissues of female anophelines that have undergone low-intensity *P. falciparum* infections representative of those occurring naturally in disease-endemic regions. To our eyes, the significant reduction in prevalence and other experimental challenges associated with these studies are outweighed by the necessity to examine laboratory-based *Plasmodium* infections under the most biologically relevant conditions possible. Malaria-infected mosquitoes along with their naive counterparts were subsequently aged, thereby facilitating the development of salivary-gland-stage sporozoites. As might be expected, these infections have broad effects on transcript abundance (as a partial proxy for differential gene expression) across a large number of gene families. Here, we have focused on those that impact neuronal function and chemosensation as well as age-related immunity and metabolic homeostasis.

The transcriptome profile shifts we have uncovered reveal a variety of systematic impacts that broadly indicate there is a synergistic advantage for mosquitoes harboring salivary gland sporozoites (and therefore likely to be malarious) that aligns with significant increases in their vectorial capacity (Figure 13). These notably include an expected increase in chemosensory sensitivity that could reasonably lead to more successful host seeking of infectious mosquitoes, along with a surprisingly distinctive transcriptome alteration that collectively aligns with an anti-aging paradigm that would likely provide a distinct advantage to these mosquitoes. When compared with similarly aged, uninfected mosquitoes, the synergistic advantage of naturally relevant malaria infections provides at least a partial rationale for the enduring persistence of *Plasmodium* pathogens and, indeed, global malaria.

**Figure 13.**
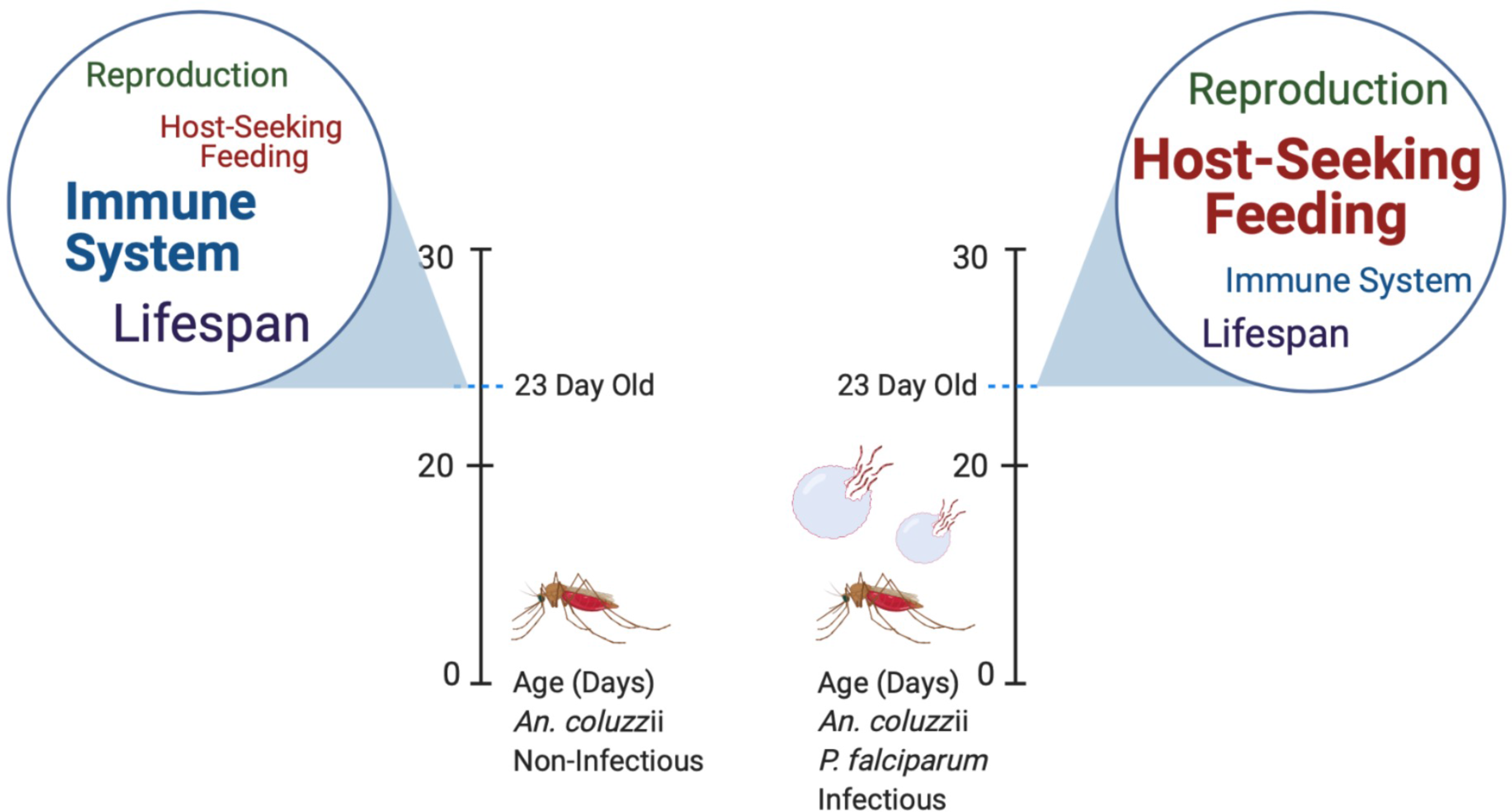
Schematic depicting transcriptional shifts in *An. gambiaei* following infection with *P. falciparum* sporozoites compared to naïve controls. Text is weighted to emphasize degree of transcript abundance change.

## Materials and Methods

### Ethics Statement

Commercial anonymous human blood provided by the Johns Hopkins Institute Core Facility, Baltimore, MD, was used for parasite cultures and mosquito feeding and as such did not require informed consent. The Johns Hopkins School of Public Health Ethics Committee approved the mosquito rearing protocols used in this study.

### Mosquito Rearing and Plasmodium Infections

*An. gambiae* s.s. (*An. gambiae* Keele strain; Ranford-Cartwright et al., 2016) mosquitoes were reared and maintained at 27 °C and 80% humidity with a 12-h light:12-h dark cycle at Johns Hopkins School of Public Health insectary (Baltimore, MD) according to standard procedures (Dong et al., 2009). Newly emerged, 0-day-old female *An. gambiae* were split into nine independent lines (groups; six infected or treatment, and three non-infected or control). All control and treatment groups were given their first and only bloodmeal at 5 days old (Haymer et al., 1997). Treatment groups were fed live *P. falciparum* NF54 wild-type (NF54W) gametocyte cultures at a final gametocytemia of 0.008% in human blood mix (provided by the Johns Hopkins Institute Core Facility, Baltimore, MD) through glass membrane feeders at 37°C for 1 h (Figure 1). Controls were fed non-infected naïve commercial anonymous human blood in parallel. All mosquitoes were starved for 3-6 h prior to maximize blood feeding and engorgement. After feeding, treatment and control lines were independently sorted at 4°C and unfed and partially fed mosquitoes were removed and disposed. The remaining fully engorged mosquitoes in treatment and control lines were incubated for 18 days in an insectary maintained at 27°C and 80% relative humidity, with a photoperiod of 12h light:12h dark, with dusk and dawn periods of 1 h each at the beginning of each scotophase. All lines had continuous access to sterile distilled water and a 10% sucrose/distilled water solution throughout the duration of the study. The infection of *P. falciparum* was confirmed at the oocyst stage by sampling at least 8-10 mosquitoes from each replicate at 8 days post infection (Table S1). In total, 58 randomly selected mosquitoes were dissected in PBS, stained with 0.1% PBS-buffered mercurochrome (MilliporeSigma, Burlington, MA) and examined under a light-contrast microscope (Olympus Life Science, Waltham, MA). At the oocyst stage, *P. falciparum* infection prevalence of the 58 randomly selected infected mosquitoes was 62%, with a median count of two oocysts per mosquito midgut (SEM = 0.33).

### Mosquito Dissections and Nucleic Acid Extractions

After collection, each mosquito in this study was individually stored prior to manual trans-section of heads (including sensory appendages) and thorax including salivary glands, referred to as ‘head’ or ‘head tissues’, from body/carcass (stored for future use). Total RNAs extracted from the heads of established treatment- and control-line *An. gambiae* mosquitoes were used for Illumina NovaSeq sequencing. Total DNAs were also extracted from the heads of treated mosquitoes for PCR to validate the *P. falciparum* infection at sporozoite stage in individual mosquitoes. All head tissues were collected from control- and treatment-line mosquitoes immediately following the 18-day incubation period (Figure 1). Dissections of control- and treatment-line mosquito heads were performed over 3 days; each line was handled on a different day to prevent cross contamination. All mosquitoes were handled wearing gloves and all dissection equipment washed with glassware detergent, autoclaved, and decontaminated with RNase Away (MilliporeSigma, Burlington, MA). To eliminate potential hemolymph contamination during mosquito dissections, all equipment was sterilized using 90% ethanol and decontaminated with RNase Away (MilliporeSigma, Burlington, MA) in between individual mosquito handling. To maintain tissue integrity, dissections were performed on cold-anesthetized mosquitoes, and work limited to one line—treatment or control. For treatment lines, the head with intact salivary glands from one mosquito was dissected into a distinct numbered and cataloged 1.5- mL test tube containing 200µL Tri Reagent (MilliporeSigma, Burlington, MA). The smaller starting volume of Tri Reagent (MilliporeSigma, Burlington, MA) aided subsequent tissue homogenization.

PCR infection validation studies on treatment-line mosquitoes required that all dissected heads remain separated. Prior to dissections, each treatment-line *An. gambiae* mosquito was randomly assigned an identification number used for labeling of all associated dissected tissues. In total, the heads of 176 treatment-line mosquitoes were collected into 176 distinct labeled tubes. For control lines, the heads from 30 mosquitoes within a single line were dissected and pooled into one labeled 1.5-mL test tube containing 400 µL Tri Reagent (MilliporeSigma, Burlington, MA). This process was repeated for the two remaining control lines. All sample tubes containing dissected head(s) in Tri Reagent (MilliporeSigma, Burlington, MA) were frozen at -70°C in preparation for RNA and DNA extractions.

For RNA extractions, sample tubes were processed in small batches of ten, with treatment and control samples handled separately to prevent cross contamination. A batch of ten sample tubes containing *An. gambiae* head tissues in Tri Reagent (MilliporeSigma, Burlington, MA) was thawed on ice and the head tissues were homogenized first manually using individual sterile, DNA/RNA-free disposable pellet pestles (Fischer Scientific, Walthom, MA) and second, mechanically by sonication (Q700; QSonica, Newtown, CT) to ensure thorough head tissue and cell disruption. Sonication was performed in short 10- to 20-s bursts with samples on ice to minimize nucleic acid shearing. To prevent cross contamination, one pellet pestle was used per sample tube, and the sonicator probe was thoroughly sterilized using 90% ethanol and decontaminated with RNase Away (MilliporeSigma, Burlington, MA) in between individual sample processing. Following complete tissue homogenization, RNA extractions were performed. An additional 200µL Tri Reagent (MilliporeSigma, Burlington, MA) was added to each treatment sample tube bringing the total volume to 400 µL, equivalent to control sample volumes. Sample tubes were then hand shaken and incubated at room temperature for 5 min. A 100-µL aliquot of UltraPure phenol:chloroform:isoamyl alcohol (25:24:1, v/v; Thermo Fischer, Walthom, MA) was added to each of the tubes, which were then vigorously hand shaken for 1 min and incubated at room temperature for 5 min. For complete phase separation, sample tubes were centrifuged at 13,000 RPM for 15min at 4°C. The resulting aqueous phase (containing RNAs) from each sample tube was carefully transferred into a coordinately labeled new RNase-free 1.5-mL test tube containing 100µL 70% ethanol and thoroughly mixed by pipetting. Sample tubes containing the remaining interphase and organic phase were frozen at -70°C in preparation for DNA extractions.

Total RNAs were extracted from the aqueous phase/ethanol solutions according to the manufacturer’s protocol using the PureLink RNA Mini Kit (Thermo Fischer, Waltham, MA). This process was repeated for all 179 treatment and control sample tubes. Total RNA concentrations and purities were measured using NanoDrop (Thermo Fischer, Waltham, MA) and any detected phenol contaminants removed by 1-butanol extraction (Krebs et al., 2009). Control-sample total RNAs were kept separate to allow for three distinct biological replicates pertinent to RNAseq analyses. For the treatment replicates, after each individual sample was confirmed to contain *P. falciparum* sporozoites via PCR validation, three distinct pools were prepared for RNAseq analyses. Ultimately, 65 of the 176 treatment samples were positively confirmed for *P. falciparum* infection and total RNAs were pooled into three replicates (21 samples per replicate).

DNA extractions were performed on all treatment sample tubes containing remaining interphase and organic phase solutions to perform PCR *P. falciparum* validation studies. Like RNA extractions, treatment samples were processed in small batches of ten to preserve DNA integrity. A batch of treatment sample tubes was thawed on ice, centrifuged at 13,000 RPM for 5 min at 4°C, and any remaining aqueous phase was removed and disposed of to prevent RNA contamination of DNA extracts. Then, 200µL back extraction buffer was added to each sample tube and hand mixed by inversion for 3 min. For complete phase separation, sample tubes were centrifuged at 13,000 RPM for 30 min at 4°C. The resulting aqueous phase (containing DNAs) from each sample was carefully transferred into a coordinately labeled, new sterile 1.5-mL test tube containing 200µL 70% ethanol and thoroughly mixed by pipetting. Total DNAs were extracted from the aqueous phase/ethanol solutions according to the manufacturer’s protocol using the DNAEasy Blood and Tissue Kit (Qiagen, Germantown, MD). Total DNA concentrations and purities were measured using NanoDrop (Thermo Fischer, Walthom, MA) and any detected phenol contaminants removed by 1-butanol extraction (Krebs et al., 2009). The total gDNA was finally resuspended at a final concentration of 10ng/µL and kept at -20°C for further PCR experiments.

### PCR-Based Plasmodium infection validation

PCR experiments were performed to validate the presence of *P. falciparum* sporozoites within each treatment sample. PCR experiments used total DNAs extracted from dissected heads and salivary glands of individual treatment-line *An. gambiae* female mosquitoes (Ref. *Mosquito Dissections and Nucleic Acid Extractions*). PCR was performed using an MJ Thermal Cycler (BioRad Inc, Hercules, CA) in combination with the DreamTaq Mastermix© technology (Thermo Scientific, Waltham, MA) according to the manufacturer’s protocol. PCR products were visualized on a 1% agarose gel using the Quick-load Purple 1-Kb DNA ladder (New England Biolabs, Ipswich, MA). Primer3 Plus (Untergasser et al., 2012) was used to design primer pairs for the *P. falciparum* circumsporozoite gene (CS) and the *An. gambiae* ribosomal s7 gene (AgRsp7) used as a reference for internal standardization (Coppi et al., 2011). The primer sets used were as follows:

CS forward (Fw): 5’-ACGAGAAATTAAGGAAACCAAAACA- 3’. CS reverse (Rv): 5’ -ACCTTGTCCATTACCTTGATTGT- 3’. *AgRsp7*Fw: 5’ -AGAATCGAACTCTGGTGGCTG- 3. *AgRsp4*Rv: 5’ -ACACAACATCGAAGGATACGA- 3’.

All PCR primers were validated and optimized in preliminary studies. PCR experiments were conducted for all treatment samples, with a correlating negative control (blank) and positive *P. falciparum* infected positive control (Figure 2). Treatment samples producing CS PCR products were reconfirmed to be positively infected through a second follow-up PCR experiment.

### Illumina NovaSeq Paired-end Sequencing

Deep sequencing utilizing Illumina RNAseq technology (Illumina, San Diego, CA) was performed at Hudson Alpha Discovery Genomic Services Laboratory (Huntsville, AL). Poly(A) mRNAs were isolated separately for library preparation and barcoding from approximately 500ng of total RNA collected from each of the three control replicates and approximately 300ng of total RNA collected from each of the three treatment replicates. Final mRNAs were fragmented, and cDNA was synthesized, amplified, digested, and purified using Illumina TruSeq chemistry protocols (Illumina, San Diego, CA). In preparation for sequencing, each of the six libraries—three treatment and three controls—was labelled with a 16-bp barcode to identify sequencing data generated from each library [treatment replicate 1 (T1) =TTGCAGGTTGCAGCGG, treatment T2 = CAACTGTATGAGTGCC, treatment T3 = AGTCAATGTAGAGGCC, control C1 = ACAAGCTTACAACTAT, control C2 = CGTCCTAACCATCGGC, and control C3 = TTGAGAACTCTCTAAC]. Barcoded treatment and control replicate libraries were hybridized across four lanes of two S4 flow cells for cBot (Illumina, San Diego, CA) cluster generation and sequencing (paired-end, 2 x 150bp; total 300 cycles). A total of 36 raw data sets of sequencing outputs, 6 per library, were generated using the Illumina software assembler (Illumina, San Diego, CA).

### Illumina Bioinformatics

The 36 control and treatment replicate Illumina NovaSeq data sets generated were cleaned and quality trimmed (Q30) using Trimmomatic v0.33 (Bolger et al., 2014), then verified for quality control using FastQC v.0.11.9 (https://www.bioinformatics.babraham.ac.uk/projects/fastqc/). Control and treatment replicate data sets were aligned to the *An. gambiae* PEST strain genome (Vectorbase- 53_AgambiaePEST_Genome.fa, https://vectorbase.org) using STAR v.2.7.5 (https://github.com/alexdobin/STAR), with *An. gambiae* PEST strain gene annotations (Vectorbase- 53_AgambiaePEST.gff, https://vectorbase.org) to map splice junctions, 2-pass mapping, -- sjdbOverhang set to 149bp, MAPQ set to 60, --alignIntronMin set to 10 bp, and a preliminary UNIX concatenate script resulting in six output BAM alignment files: control R1, control R2, control R3, treatment R1, treatment R2, and treatment R3. HTSeq v0.11.2 (Anders et al., 2015), an alignment-based count algorithm, was used to quantify transcript counts for all six BAM alignment files with default parameters and mapping validation. DESeq2 (Bioconductor v.3.8) and R v.3.5.1 were used to calculate mean variance and obtain differentially abundant transcripts between control and treatment replicates using default parameters with added shrinkage and Wald’s hypothesis tests. Gene ontologies (GO) of differentially abundant transcripts were obtained from OmicsBox using the GenBank non-redundant protein (nr) database containing all available mosquito genomes with an expected value (e-value) cutoff of less than 0.01. All GO annotations were thereafter manually confirmed via BLAST, and phylogenetic analyses and FPKM values for differentially abundant transcripts were analyzed for significance using ANOVA with Sidak’s multiple comparison test using Prism™ v.8 (GraphPad, La Jolla, CA).

## Acknowledgments

We thank Ms. Maureen Ubani for technical assistance for PCR validations; Dr. Antonis Rokas, Dr. H. W. Honegger and Dr. Stephen Ferguson for their comments on this manuscript as well as other members of the Zwiebel lab for critical suggestions during the course of this work. We also thank Dr. A.M. McAinsh for editorial assistance as well as the Johns Hopkins Malaria Research Institute insectary and parasitology core facility, for mosquito rearing and other technical help. This work was conducted with the support of the Bloomberg Philanthropies (to G.D.), Vanderbilt University (to LJZ) and National Institutes of Health grants (NIAID, AI122743) to GD, and (NIAID, AI127693) to LJZ.

## Author Contributions

A.L.C. and L.J.Z. designed the study. A.L.C. and Y. D. performed the experiments. A.L.C and D.R. analyzed data and prepared figures; A.L.C, D.R., Y.D., G.D. and L.J.Z. interpreted the results and wrote the manuscript.

---

**Figure S1.**
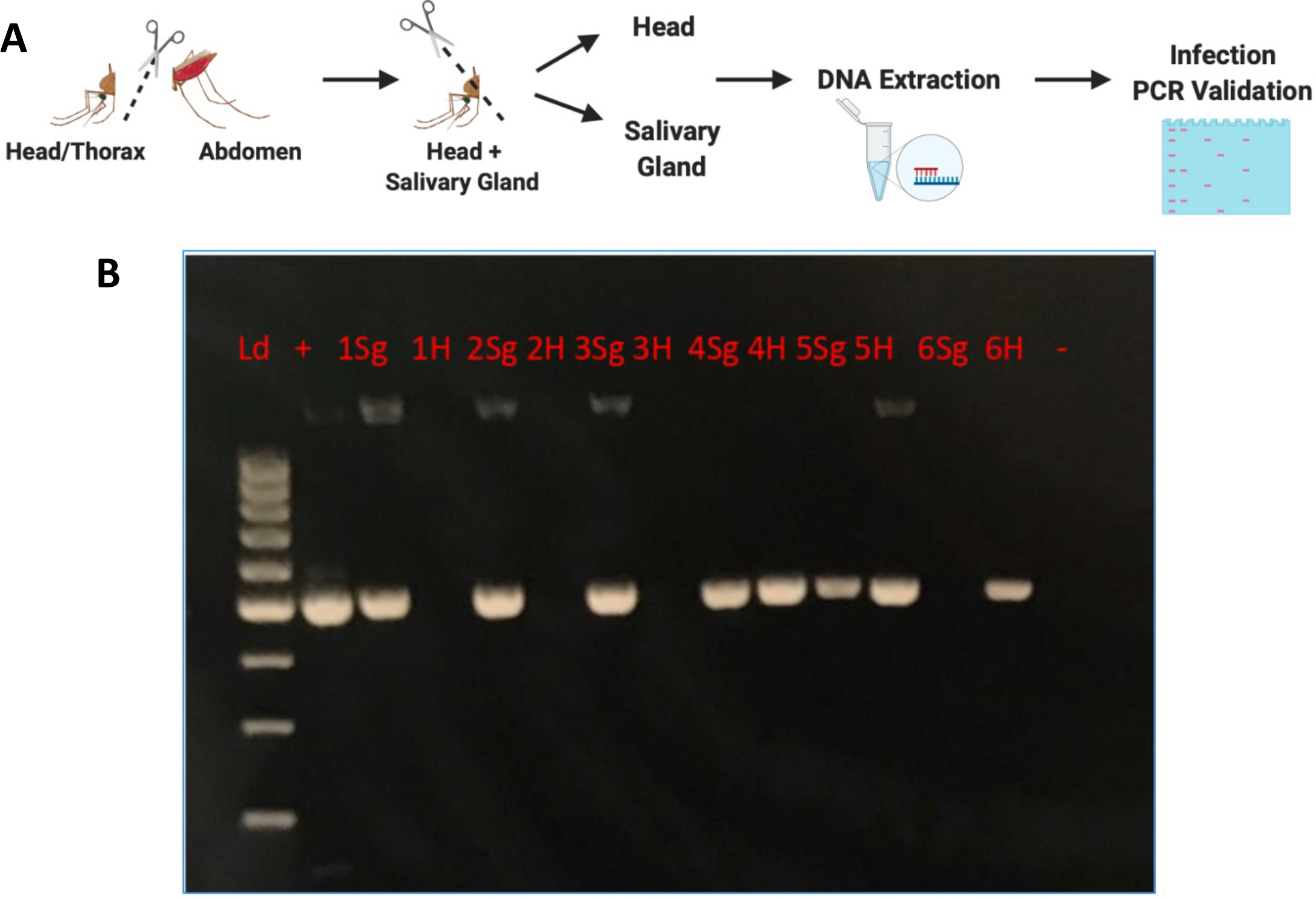
(A) Schematic representation of gDNA extraction methodology from head and salivary gland tissues of *P. falciparum* sporozoite infected treatment mosquitoes. (B) Agarose gel (1.5%) electrophoresis image of amplified products using CS2 primer sets. Lanes 1Sg and 1H represent examined salivary gland and head tissues, respectively from treatment *An. gambiae* s.s. (*An. gambiae*) mosquito 1. Remaining lanes follow a like organization for 5 additional treatment *An. gambiae* mosquitoes with a (Kazuya Ujihara) positive and (-) negative control. Lane Ld, 100bp DNA size marker.

